# Stomatal sensitivity to VPD across 38 tree species reflects past drought responses but is little explained by stomatal and leaf economic traits

**DOI:** 10.64898/2026.09.09.750341

**Authors:** Lena Sachsenmaier, Ronny Richter, Camilla Ahner, Lena Kretz, Anja Kahl, David Schellenberger Costa, Florian Schnabel, Ingmar Staude, Manon Sabot, Christian Wirth

**Affiliations:** German Centre for Integrative Biodiversity Research (iDiv) Halle-Jena-Leipzig, Leipzig, Germany; Systematic Botany and Functional Biodiversity, Institute of Biology, Leipzig University, Leipzig, Germany; Chair of Silviculture, University of Freiburg, Freiburg, Germany; Future Forests Cluster of Excellence, University of Freiburg, Freiburg, Germany; Max-Planck Institute for Biogeochemistry, Jena, Germany

**Keywords:** climate change, drought, stomatal behaviour, stomatal conductance (g_s_), vapour pressure deficit, leaf traits, tree water-use, δ^13^C

## Abstract

Climate change is exposing trees to rising atmospheric water demand. Because stomata regulate plant water loss, understanding stomatal behaviour is critical. However, few comparative studies examine species-specific stomatal responses to atmospheric demand.

Here, we quantify stomatal behaviour in 38 temperate tree species (27 broadleaved, 11 conifers) in a German research arboretum during a warm summer without soil drought. We measured the diurnal stomatal conductance (g_s_) of 787 leaves and modelled g_s_ species-specific responses to leaf-to-air vapour pressure deficit (VPD_L_). Four descriptors of the g_s_-VPD_L_ relationship were derived: maximum stomatal conductance (g_max_), the VPD_L_ that maximises g_s_, the VPD_L_ associated with a 50% reduction in g_s_, and the stomatal closure rate. We relate these metrics to drought-induced shifts of carbon isotope ratios and to leaf functional traits.

Large variation in stomatal behaviour exists, ranging from early-closure to VPD_L_-tolerant strategies. Species maintaining g_s_ at higher VPD_L_ also show weaker physiological responses to past drought. g_max_ is partly explained by leaf economic traits; other gs-VPDL descriptors are not related to stomatal anatomy and leaf economy.

Our interspecific comparison reveals substantial variation in stomatal responses to atmospheric demand that is relevant to drought responses, but cannot be readily inferred from commonly measured leaf traits.

## Introduction

Climate change is leading to an increase in the frequency and intensity of concurrent drought and heat waves, with profound consequences for forest health and functioning worldwide (Gazol & Camarero, 2022; Hari et al., 2020; Markonis et al., 2021; Werner et al., 2026). While drought impacts on trees have traditionally been interpreted in terms of limited soil moisture, atmospheric dryness, quantified as vapour pressure deficit (VPD), can be an additional driver of physiological stress (Grossiord et al., 2020; Novick et al., 2024). Elevated temperatures can increase VPD, thereby forcing trees to regulate water loss under increased transpirative (atmospheric) demand, independently of soil moisture. Although stomata play a central role in regulating leaf–atmosphere coupling, how stomatal conductance responds to high atmospheric demand remains insufficiently characterised.

Stomatal control strategies are often characterised using a relatively small number of tree species (e.g., Fernández-Molano et al., 2025; Sachsenmaier et al., 2025; Tsuji et al., 2020), with equivocal evidence for whether observed patterns generalise across taxa (Buckley, 2019). Existing studies suggest substantial diversity in stomatal responses to VPD among species with contrasting functional characteristics (e.g. Kröber & Bruelheide, 2014), calling for a broad examination of species to identify generalisable patterns of stomatal behaviour, i.e., variation in the magnitude of stomatal conductance and in how stomata regulate it. Understanding such patterns could help better integrating stomatal behaviour into trait-based approaches and ecosystem models. Because direct measurements of stomatal responses to VPD are resource-intensive, linking this behaviour to widely measured functional traits of leaf anatomy and economy would offer an efficient way to extend predictions of stomatal behaviour across diverse tree species.

Stomata regulate gas exchange between the leaf interior and the atmosphere, balancing CO_2_ uptake for photosynthetic carbon assimilation against water loss through transpiration (Cowan & Farquhar, 1977). Stomatal conductance (g_s_) quantifies the ease with which CO_2_ or water vapour diffuses through stomatal pores (Farquhar & Sharkey, 1982); hereafter, g_s_ refers to conductance to water vapour from the leaf interior outward. Stomata open in response to light to allow CO₂ into the leaf for photosynthesis, but this inevitably increases water loss as vapour diffuses out to the drier atmosphere. Stomata therefore close when the benefit of continued CO₂ uptake declines or when the risk of water loss increases, such as under low light towards the end of the diel cycle, in response to soil water limitation, or/and high VPD. Stomata typically respond to changes in VPD within minutes (Kaiser & Kappen, 2001; McAdam & Brodribb, 2015; Vialet-Chabrand et al., 2017), as stomatal aperture is regulated by changes in guard-cell turgor resulting from coordinated hydraulic and biochemical signals. Changes in leaf water status can alter guard-cell turgor directly, while chemical signals such as abscisic acid (ABA) can modify guard-cell ion transport, water fluxes, and thereby stomatal aperture (Buckley, 2005, 2019; Grossiord et al., 2020; McAdam & Brodribb, 2016).

The sensitivity, speed, and magnitude of stomatal regulation in response to hydraulic and environmental signals, to which we hereafter refer collectively as ‘stomatal sensitivity’, vary considerably among species (Lawson & Blatt, 2014; McAusland et al., 2016), yet no widely agreed-upon framework exists for quantifying these responses across tree species. Much of the literature has focused on stomatal closure in relation to soil water stress and leaf water potential (e.g., Brodribb & Holbrook, 2003; Henry et al., 2019; Klein, 2014; Martin-StPaul et al., 2017; Waite et al., 2024); fewer studies have characterized differences in species’ responses to VPD in isolation (e.g., Cunningham, 2004; Gao et al., 2015; Kröber & Bruelheide, 2014; Oren et al., 1999). Importantly, species operate over different ranges of leaf water potential, such that stomatal sensitivity to declining leaf water potential does not necessarily indicate when stomata will close under increasingly dry conditions (Martínez-Vilalta & Garcia-Forner, 2017). Plant trait databases such as TRY (Kattge et al., 2020) often only store information on a few descriptors of stomatal behaviour, e.g., the maximum, mean, and minimum stomatal conductance, which describe magnitudes under specific conditions but not its regulation. Because stomatal conductance is highly dynamic, comparisons based on measurements taken under different environmental conditions may confound species differences with environmental effects (Miner et al., 2017). Measuring many functionally divergent species under comparable conditions therefore appears a necessary requisite for characterizing interspecific variation in stomatal behaviour.

Knowing the full extent of variation in stomatal control may be important as it has been shown to be relevant to tree growth during drought in subtropical and temperate species (Sachsenmaier et al., 2025; Schnabel et al., 2021, 2024), likely owing to the stomata in balancing carbon gain with the risk of hydraulic dysfunction. If variation in stomatal behaviour consistently reflects species strategies in dealing with heat and drought stress, the physiological consequences of these strategies may therefore also be detectable in integrated responses to a drought event. As atmospheric demand rises and soil water becomes limiting, stomatal closure becomes the primary mechanism by which trees limit water loss and avoid hydraulic failure, making it a central component of the drought response (Martínez-Vilalta & Garcia-Forner, 2017).

To understand and validate the interspecific diversity of stomatal behaviour, leaf carbon isotope discrimination (δ^13^C) can be employed. δ^13^C integrates the balance between carbon assimilation and water loss over time and is widely used as an indicator of plant water use efficiency (Cernusak et al., 2013; Farquhar et al., 1989; Goldsmith et al., 2023). Reductions in stomatal conductance decrease intercellular CO_2_ concentrations thereby reducing discrimination against the heavier carbon isotope (^13^C). Plants that kept their stomata closed for longer periods would generally show less negative (higher) δ^13^C values. Importantly, severe drought events typically involve multiple co-occurring stressors, including high temperature and atmospheric water demand alongside reduced soil water availability. Comparisons of a tree’s δ¹³C across contrasting water availability conditions in the field (expressed as the difference in δ¹³C between dry and wet years) therefore captures integrated drought-induced physiological responses, including those associated with stomatal regulation (Jing et al., 2024; Jucker et al., 2017; Schnabel et al., 2022). Yet, whether species differences in baseline stomatal behaviour reflect such integrated drought-induced changes in leaf δ^13^C remains to be explored. Identifying leaf functional traits associated with the different dimensions of stomatal behaviour could then help mechanistically explain and generalize patterns of stomatal behaviour across species.

Anatomical stomatal traits are likely to be predictors of the magnitude of stomatal conductance. Stomatal density (SD) and stomatal size (guard cell length, GCL) have been identified as key factors influencing the anatomical conditions for maximum stomatal conductance (Hetherington & Woodward, 2003). Together, they affect the total surface area over which gas exchange occurs, and thus largely determine the maximum pore area available for diffusion (besides pore depth). Because smaller pores have a shorter diffusion path, for a given stomatal area, many small stomata typically achieve a greater g_max_ than a few large stomata (Franks & Beerling, 2009); this helps to explain the generally higher g_max_ of angiosperms compared to gymnosperms (de Boer et al., 2016). Stomatal anatomy may also influence stomatal sensitivity. Smaller stomata could confer an advantage owing to faster opening and closing (Drake et al., 2013; Hetherington & Woodward, 2003; Raven, 2014): if species with a high density of smaller stomata can close more quickly and with greater precision, this would allow for a better utilization of favourable photosynthetic conditions (Drake et al., 2013; Lawson & Blatt, 2014). Those species may also tolerate relatively high VPD for longer as they could either reduce conductance rapidly once atmospheric demand increases beyond their comfort zone, or maintain partial stomatal opening in leaf regions where conditions are still favourable, e.g., near major veins (Mott & Buckley, 1998, 2000).

Beyond stomatal anatomy, leaf traits related to the leaf economics spectrum (LES) (Wright et al., 2004) may reveal carbon uptake strategies through stomatal behaviour. Regarding the magnitude of stomatal conductance, leaf economic traits may reflect the demand side of stomatal behaviour. Species with high leaf nitrogen and specific leaf area (SLA) generally have greater photosynthetic capacity and resource acquisition rates, potentially creating a greater demand for CO_2_ and thus higher stomatal conductance, whereas species with high leaf dry matter content (LDMC) tend to have more conservative resource-use strategies (Meinzer, 2003; Onoda et al., 2017; Reich, 2014; Wright et al., 2004). Whether differences along the fast-slow investment spectrum, pronounced between broadleaved and conifer species (Sanaei et al., 2026), translate into distinct stomatal closure behaviour remains unclear. On the one hand, conservative species tend to also be more conservative in their overall water-use strategy, and may be expected to close earlier to protect hydraulic functions (Skelton et al., 2015). On the other hand, their greater structural investments and potentially higher stress tolerance may enable them to maintain gas exchange under more negative water potentials, resulting in delayed stomatal closure. Structural traits like leaf thickness and toughness further reflect investment in mechanically robust tissues (Onoda et al., 2011), which may not only matter for withstanding the high tension associated with water stress, but also for water storage capacity, the latter which buffers rapid changes in leaf water status (Fu et al., 2019; Martins et al., 2016). Taken together, the extent to which anatomical and economic leaf traits are associated with stomatal responses to rising VPD, and whether they can explain species differences in stomatal behaviour, remains to be established.

Here, we investigate the species-specific stomatal regulation of 38 tree species (27 broadleaved, 11 conifer species) growing under common environmental conditions at a research arboretum in Germany. We derive stomatal behaviour metrics from g_s_-VPD_L_ curves modelled within a Bayesian framework based on diurnal data collected over multiple days, thus allowing for the thorough examination of species-specific responses. We then use these metrics to assess the extent to which stomatal behaviour under non-limiting soil water conditions echoes physiological drought responses as well as commonly measured leaf functional traits. We hypothesize that: (1) species differ in their stomatal behaviour strategies, with conifers showing generally lower stomatal conductance and stomatal closure at lower VPD_L_ than broadleaved species; (2) species maintaining stomatal conductance at higher VPD_L_ show smaller drought-induced shifts in leaf δ^13^C, pointing to little integrated physiological adjustments to drought; (3) species with a high density of small stomata exhibit higher stomatal conductance but close their stomata at a higher VPD_L_; (4) species with conservative leaf economic traits display lower stomatal conductance and closure at higher VPD_L_ than species with acquisitive leaf traits.

## Materials and Methods

### Study Area and Experimental Design

The study was conducted at the ARBOfun research arboretum, located in Großpösna, south-east of Leipzig, Germany (51°16′N, 12°30′E; 150 m a.s.l.). Established between 2012 and 2014 on 2.5 hectares of former agricultural land, the site hosts ca. 100 tree species spanning 39 families. Trees were planted following a randomized block design, with five blocks comprising one individual per species, and a wide spacing of 5.8 m between trees. The site has a temperate climate with a mean annual temperature of 9.7 °C and mean annual precipitation of 520 mm (DWD Climate Data Center, Station Leipzig/Halle, ID 2932). The soil is classified as Luvisol (Ferlian et al., 2017). For the present study, a subset of 38 functionally divergent tree species (27 angiosperms, 11 conifers) was selected (see Table 1).

**Table 1:** Tree species included in the study, grouped by major clade (27 angiosperms, 11 gymnosperms), along with their common English names and plant family. Please note that scientific names follow the current status of the World Checklist of Vascular Plants (WCVP) and names commonly used in previous literature are provided in parentheses.

| Clade | Species code | Species | Common name | Family |
| --- | --- | --- | --- | --- |
| Angiosperm | Ace_neg | *Acer negundo* L. | Boxelder maple | Sapindaceae |
|  | Ace_pla | *Acer platanoides* L. | Norway maple | Sapindaceae |
|  | Ace_pse | *Acer pseudoplatanus* L. | Sycamore maple | Sapindaceae |
|  | Aes_hip | *Aesculus hippocastanum* L. | Horse chestnut | Sapindaceae |
|  | Aln_glu | *Alnus glutinosa* (L.) Gaertn. | Black alder | Betulaceae |
|  | Bet_pen | *Betula pendula* Roth | Silver birch | Betulaceae |
|  | Car_bet | *Carpinus betulus* L. | European hornbeam | Betulaceae |
|  | Cas_sat | *Castanea sativa* Mill. | Sweet chestnut | Fagaceae |
|  | Cor_dom | *Cormus domestica* (L.) Spach<br>(previously: *Sorbus domestica*) | Service tree | Rosaceae |
|  | Cor_col | *Corylus colurna* L. | Turkish hazel | Betulaceae |
|  | Fag_syl | *Fagus sylvatica* L. | European beech | Fagaceae |
|  | Fra_exc | *Fraxinus excelsior* L. | European ash | Oleaceae |
|  | Jug_reg | *Juglans regia* L. | English walnut | Juglandaceae |
|  | Pla_his | *Platanus x hispanica* Mill. Ex Münchh.<br>(previously: *Platanus x acerifolia*) | London plane | Platanaceae |
|  | Pop_tre | *Populus tremula* L. | European aspen | Salicaceae |
|  | Pru_avi | *Prunus avium* (L.) L. | Wild cherry | Rosaceae |
|  | Que_pub | *Quercus pubescens* Willd. | Pubescent oak | Fagaceae |
|  | Que_rob | *Quercus robur* L. | Pedunculate oak | Fagaceae |
|  | Rob_pse | *Robinia pseudoacacia* L. | Black locust | Fabaceae |
|  | Sal_cap | *Salix caprea* L. | Goat willow | Salicaceae |
|  | Sal_fra | *Salix fragilis* L. | Crack willow | Salicaceae |
|  | Sor_auc | *Sorbus aucuparia* L. | Rowan | Rosaceae |
|  | Til_cor | *Tilia cordata* Mill. | Small-leaved lime | Malvaceae |
|  | Til_tom | *Tilia tomentosa* Moench | Silver lime | Malvaceae |
|  | Tor_gla | *Torminalis glaberrima* (Gand.) Sennikov & Kurtto<br>(previously: *Sorbus torminalis*) | Wild service tree | Rosaceae |
|  | Ulm_lae | *Ulmus laevis* Pall. | European white elm | Ulmaceae |
|  | Ulm_min | *Ulmus minor* Mill. | Field elm | Ulmaceae |
| **Gymnosperm** | Abi_alb | *Abies alba* Mill. | European silver fir | Pinaceae |
|  | Abi_gra | *Abies grandis* (Douglas ex D.Don) Lindl. | Grand fir | Pinaceae |
|  | Ced_deo | *Cedrus deodara* (Roxb. ex D.Don) G.Don | Deodar cedar | Pinaceae |
|  | Lar_dec | *Larix decidua* Mill. | European larch | Pinaceae |
|  | Pic_abi | *Picea abies* (L.) H.Karst. | Norway spruce | Pinaceae |
|  | Pic_sit | *Picea sitchensis* (Bong.) Carrière | Sitka spruce | Pinaceae |
|  | Pin_nig | *Pinus nigra* J.F.Arnold | Black pine | Pinaceae |
|  | Pin_pon | *Pinus ponderosa* Douglas ex. C.Lawson | Ponderosa pine | Pinaceae |
|  | Pin_syl | *Pinus sylvestris* L. | Scots pine | Pinaceae |
|  | Pse_men | *Pseudotsuga menziesii* (Mirb.) Franco | Douglas-fir | Pinaceae |
|  | Tax_bac | *Taxus baccata* L. | European yew | Taxaceae |

### Stomatal Conductance Measurements

We measured stomatal conductance (g_s_) across the selected species. For each species, we measured seven mature, undamaged leaves in three individuals (787 leaves, 114 trees in total); all leaves were sun-exposed, from the upper third of the canopy, and on different but south-east oriented branches. We marked the leaves to ensure repeated measurements throughout the day. For compound-leaved angiosperms, measurements were consistently taken on the terminal leaflet. For gymnosperms, measurements were taken from the same needle bundle (*Pinus* spp.) or branchlet, using last year’s shoots to ensure needle maturity (except for deciduous *Larix decidua*).

Across eight days in June and July 2024, we measured g_s_ at intervals of 45-90 min, between 6:00 AM and 4:00 PM, capturing a large part of the diurnal cycle. Data collection was restricted to mostly clear-sky conditions and low wind speeds (Table S1). While mean air temperature and air vapour pressure deficit (VPD) showed variation across the eight sampling days (Table S1), air VPD reached values of up to 2-3 kPa in the afternoons, values large enough to induce stomatal closure at midday and, thus, for the later analysis based on g_s_-VPD_L_ curves (Figure S1). Light availability varied diurnally and was therefore accounted for as a covariate in our analyses. Although soil moisture declined over the course of the sampling campaign (Figure S2b), the system remained outside of drought conditions (Figure S2a), such that soil water stress effects on physiological behaviour were deemed negligible. Further, by systematically stratifying the three individuals of each species across different measurement days, we minimized confounding temporal effects (Figure S3).

We measured g_s_ with steady-state handheld porometers - LI-600 for broadleaves and LI-600N for needle-leaved species (LI-COR, Lincoln, Nebraska, USA). Because the instruments integrate different chamber designs for broadleaved and needle-leaved foliage, respectively, g_s_ per unit leaf area reflects the abaxial surface for angiosperms and the total projected surface area for gymnosperms. For gymnosperms, instrument settings and measurements were adapted to account for species-specific needle dimensions (Table S2; Note S1). To prevent sensor drift, automatic relative humidity (RH) sensor matching was performed with an empty chamber prior to measuring each new tree. All measurements underwent multi-stage quality control to ensure their physical and physiological plausibility (see Note S2). The final dataset comprised 6002 single observations from 38 species, from 18 to 21 leaves per species and at least six measurements per leaf across a measurement day.

### Modeling stomatal conductance as a function of VPDL

Since stomata respond to leaf-level atmospheric conditions, we used leaf-to-air VPD (VPD_L_) measured by the porometer for modelling, which ensures a direct temporal match between g_s_ and the microclimatic conditions experienced by the leaf. We modelled the effect of VPD_L_ on stomatal conductance (g_s_) as:

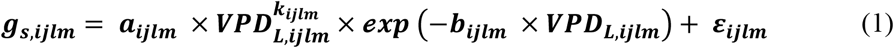

where *i, j, l,* and *m* denote species, tree, leaf and observation, respectively. Parameter *a* controls the overall magnitude of the response, *k* the initial increase of g_s_ with low VPD_L_, and *b* its decline at high VPD_L_. The parameter submodels were:

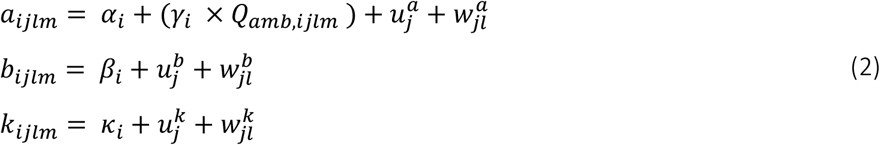

where Q_amb_ is the centred and scaled ambient light associated with each observation, *α_i_*, *β_i_* and *κ_i_* are species-specific effects, and γ_i_ represents the species-specific effect of ambient light on α. The terms *u* and *w* represent tree- and leaf-level random effects, respectively (Figure 1).

**Figure 1:**
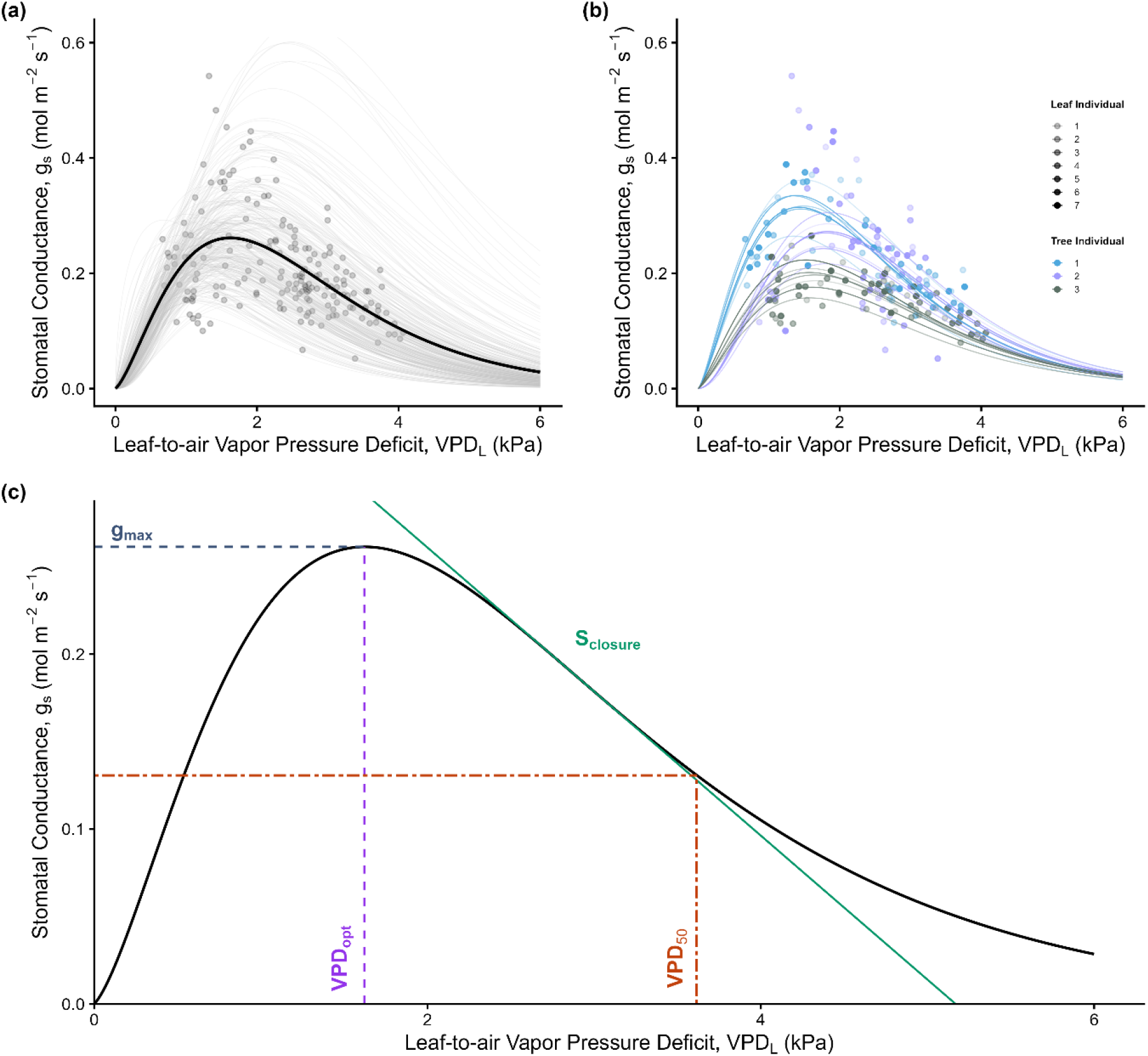
Example Bayesian hierarchical model fits of stomatal conductance (g_s_) response to leaf-to-air vapor pressure deficit (VPD_L_), for Sorbus aucuparia, with a conceptual illustration of the derived stomatal behaviour metrics. (a) Population-level relationship representing the global model structure. Points indicate the direct observations (n=181); grey lines display the posterior distribution illustrated by 300 random draws from the model parameter space, capturing uncertainty. The black line marks the mean population curve under standardized ambient light conditions (mean Q_amb_). (b) Individual-level hierarchical breakdown showing random effects partitioning. Points and modelled trajectory lines are color-coded by tree individual (n=3), and the transparency gradient distinguishes nested individual leaves within each tree (n=7). All trajectories represent the mean expected values calculated for specific leaf-level environments under standardized ambient light conditions. (c) Conceptual illustration of the stomatal behaviour metrics based on stomatal conductance (g_s_) responses to leaf-to-air vapour pressure deficit (VPD_L_). The solid line represents the reconstructed mean response curve. The maximum operational conductance (g_max_), indicated by the marine horizontal dashed line, identifies the absolute peak value of the curve and represents the maximum operational gas exchange rate captured. The corresponding optimum vapor pressure deficit (VPD_opt_), shown by the vertical purple dashed line, denotes the atmospheric demand at which g_max_ is achieved. The VPD at half-closure to a 50 % reduction in conductance (VPD_50_), marked by the orange dash-dotted line, identifies the point along the post-peak declining limb where g_s_ drops to exactly half (0.5⋅g_max)_ of its maximum operational rate. Finally, the maximum relative rate of stomatal closure (S_closure_), illustrated by the green solid tangent line, is defined as the minimum of the first derivative evaluated along the post-peak smoothed declining limb of the relative conductance curve.

Compared to not accounting for the effect of light on g_s_, our simple inclusion of a Q_amb_ effect on *a* substantially improved the model’s predictive accuracy (Δ elpd (expected log predictive density) = 428.4 ± 50.3) and further explained variance (Δ Bayesian R^2^ = 0.02). More complex formulations, including light-saturating response functions and effects of light on all three parameters (*a, b, k*), were explored but either proved unstable or induced too much equifinality.

We used a Bayesian approach to jointly estimate the nonlinear parameters, improve their identifiability and propagate parameter uncertainty through the subsequent predictions. All parameters of the non-linear function (*a, b, k*) were positively constrained using weakly informative lognormal priors, based on biologically plausible values (Table S3). Residual variation was modeled using a Student-t likelihood, with priors on the scale parameter (σ) and degrees of freedom (ν) to allow for heavy-tailed errors and robustness to outliers. Prior predictive simulations confirmed that the priors generated plausible response curves before fitting the model (Figure S4).

The model was fitted using four Hamiltonian Monte Carlo chains of 4,000 iterations (including 2,000 for warmup) with the No-U-Turn Sampler (NUTS) algorithm, as implemented in the *brms* package (Bürkner, 2018) in R (R Core Team, 2026). Convergence was assessed using the potential scale reduction factor (*R̂* < 1.01) and effective sample sizes, and posterior predictive checks were used to evaluate model performance (Tables S4, S5). Species-specific raw data and model fits are provided in Figure S5.

### Stomatal behaviour metrics

Using our fitted Bayesian model, we generated posterior predictive draws of g_s_ across a VPD_L_ vector, spanning 0–15 kPa for each species, using population-level effects only, i.e., excluding tree- and leaf-level random effects, and keeping ambient light at its mean level. All posterior draws (4000 per species) were used to propagate uncertainty in response curves and derived metrics. For each draw, species-specific response curves were constructed, smoothed using a 5-point rolling mean to reduce algorithmic noise in derivatives, and a comprehensive set of curve-based metrics was extracted (see Table 2).

**Table 2:** Stomatal behaviour metrics derived from g_s_-VPD_L_ posterior curves

| Metric | Short name | Definition | Interpretation |
| --- | --- | --- | --- |
| $g_{max}$ | Maximum $g_s$ | The peak value of the $g_s$-$VPD_L$ posterior curves. | Maximum operational gas exchange rate, measured *in-situ* |
| $VPD_{opt}$ | VPD optimum | $VPD_L$ at which $g_{max}$ is achieved. | Atmospheric demand under which stomata operate at maximum evaporative capacity (“peak comfort range”) |
| $VPD_{50}$ | VPD at half-closure | $VPD_L$ at which $g_s$ declined to 50 % of $g_{max}$ | Atmospheric demand up to which stomata operate comfortably, signalling the onset of significant $VPD_L$ stress |
| $S_{closure}$ | Closure rate | Steepest post-peak decline slope, standardized by $g_{max}$ | Relative speed of stomatal closure with increasing $VPD_L$ |

Because the response curves were reconstructed along a discrete 1000-point vector, we calculated all metrics by numerical approximation. We identified g_max_ as the peak value of the g_s_-VPD_L_ curve, and VPD_opt_ as the VPD_L_ value corresponding to this peak. Importantly, as these values are derived from field-measured data, they are inherently influenced by ambient environmental conditions, and thus represent operational rather than absolute physiological maxima. We calculated VPD_50_ as the VPD_L_ value corresponding to a 50% loss of conductance from the peak of the curve (i.e., 0.5 * g_max_). Finally, we determined the maximum rate of stomatal decline (S_closure_) by evaluating the steepest first derivative of the smoothed curve along the post-peak declining limb. Stomatal behaviour metrics were summarised at the species level using posterior means and 95% credible intervals (Table S6).

To test whether gymnosperm and angiosperm species differ in those stomatal behaviour metrics, we calculated the mean of each metric for each clade for every posterior draw and calculated the difference between gymnosperms and angiosperms. We summarized the posterior distribution of these differences using the mean and 95 % credible interval. To assess whether the four stomatal metrics captured distinct or overlapping dimensions of stomatal behaviour, we calculated pairwise Pearson correlations among all metrics and performed a principal component analysis (PCA) on the standardized species-level mean metrics.

### Leaf Trait Sampling and Measurement

#### Stomatal Density and Size

To quantify stomatal density (SD) and guard cell length (GCL) across tree species, we sampled three mature, healthy sun-exposed leaves per tree from the upper third of the canopy for each tree for which g_s_ was measured. Sampling was conducted in June 2023 for broadleaved species and in July 2025 for conifers. SD and GCL were quantified from stomatal imprints, with SD calculated per unit abaxial leaf surface for broadleaved and per unit total projected needle surface for conifers (details in Note S3 and Sachsenmaier et al. (2026a) for conifers).

#### Leaf morphological and physiological traits

Other leaf morphological and physiological traits were measured in June 2023 following standard protocols (Pérez-Harguindeguy et al., 2013). For the broadleaved trees, 10 fully expanded sun leaves were sampled per individual, with subsets used for measurements of leaf toughness, thickness, SLA, LDMC, and leaf nitrogen content per area (leaf N). For the conifers, up to 40 needles per individual were collected to obtain sufficient material. We determined the fresh mass of leaf samples, scanned the leaf area (by Expression 11000XL, Epson, UK; 1200 dpi, 24-bit colour) and measured dry mass after oven drying (70°C for 24-48 h). SLA was calculated as leaf area divided by leaf dry mass, and LDMC as leaf dry mass divided by leaf fresh mass. Leaf nitrogen content was analysed from the dried (60°C for 12 h) and homogenised material using an elemental analyser (VarioEL II, Elementar). Leaf toughness was determined using a resistance-to-punch test (Electric Test Stand TVM-N with dynamometer FH50, Sauter GmbH, Germany), and thickness was measured adjacent to the punch location with a digital caliper.

### Leaf carbon isotopes ratios

To quantify a tree’s capacity to adjust its intrinsic water-use-efficiency (iWUE), we calculated differences in leaf carbon isotope ratios between two years with contrasting water availability (Δδ^13^C; drought identification by the SPEI drought index, Figure S6). At the end of the growing season, sun leaves were collected from the same three individuals per species of the g_s_ measurements when possible. Leaves were sampled across the upper crown, dried (70°C for 48 h), ground, and analysed for their leaf carbon isotope ratio (δ^13^C, ‰) using an elemental analyser connected to an isotope ratio mass spectrometer (IRMS; BGC-IsoLab, Max Planck Institute for Biogeochemistry, Jena, Germany). δ^13^C values were calculated relative to the international VPDB standard, serving as a proxy of intrinsic water-use efficiency (Farquhar et al., 1989):

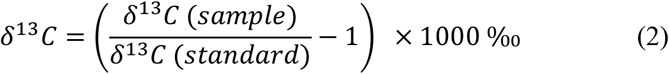

We calculated the difference in δ^13^C between the drought year (2018) and the wet reference year (2021) as

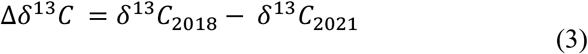

More positive values of Δδ^13^C indicate greater ^13^C enrichment during the drought year relative to the reference year and thus a greater adjustment of iWUE in response to drought, which we interpret as a greater integrated physiological drought response.

### Statistical analyses of relationships to leaf traits and Δδ^13^C

Relationships between leaf functional traits and stomatal behaviour metrics were examined via a series of Bayesian linear models using the R package *brms* (Bürkner, 2018). For each combination of stomatal behaviour metric (g_max_, VPD_opt_, VPD_50_, S_closure_; 4 metrics) and leaf trait (SD, GCL, SLA, LDMC, leaf thickness, leaf toughness, leaf N (area); 7 traits), we fitted a separate model (28 models in total).

Trait values were aggregated to species means (Table S7) and z-scored across all species. Stomatal behaviour metrics represent species-level posterior means derived from the g_s_-VPD_L_ model, with posterior standard deviations propagated as measurement uncertainty using the se() syntax in *brms*, i.e., species with greater uncertainty in their stomatal behaviour estimates contribute less information to the slope estimates proportionally. Because broadleaved and conifer species differ in their stomatal, leaf morphology and structure, tree group (broadleaved vs. conifer) was included to account for potential differences. Therefore, each model included the scaled trait, tree group, and their interaction as predictors:

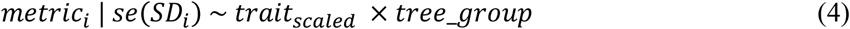

All models were fitted with four chains of 4000 iterations (2000 warmup). Group-specific slope estimates and their 95 % credible interval were extracted from the posterior distributions using the *emtrends()* function from the *emmeans* package (Lenth & Piaskowski, 2026), via the *tidybayes* interface (Kay, 2024). Model convergences were assessed via the potential scale reduction factor (*R̂* < 1.01) and effective sample size (Table S8).

To examine whether stomatal behaviour metrics were related to physiological drought responses during the past drought year 2018, we fitted analogous Bayesian linear models using species mean leaf Δδ^13^C_2018-2021_ (z-scored across species) as predictors. Uncertainty in stomatal behaviour metrics was propagated as above, and tree group was included as an interacting predictor. Models were fitted with four chains of 4000 iterations (2000 warmup) and posterior slopes were extracted as described above (Table S9).

## Results

### Stomatal conductance and stomatal behavioural spectrum

Based on repeated g_s_ measurements across a natural VPD_L_ gradient for 38 tree species, the modelled g_s_ responses to VPD_L_ (Bayesian cond. R^2^ = 0.87; Table S4, S5) revealed substantial interspecific variation in both magnitude and the shape of the response curves (Figure 2).

**Figure 2:**
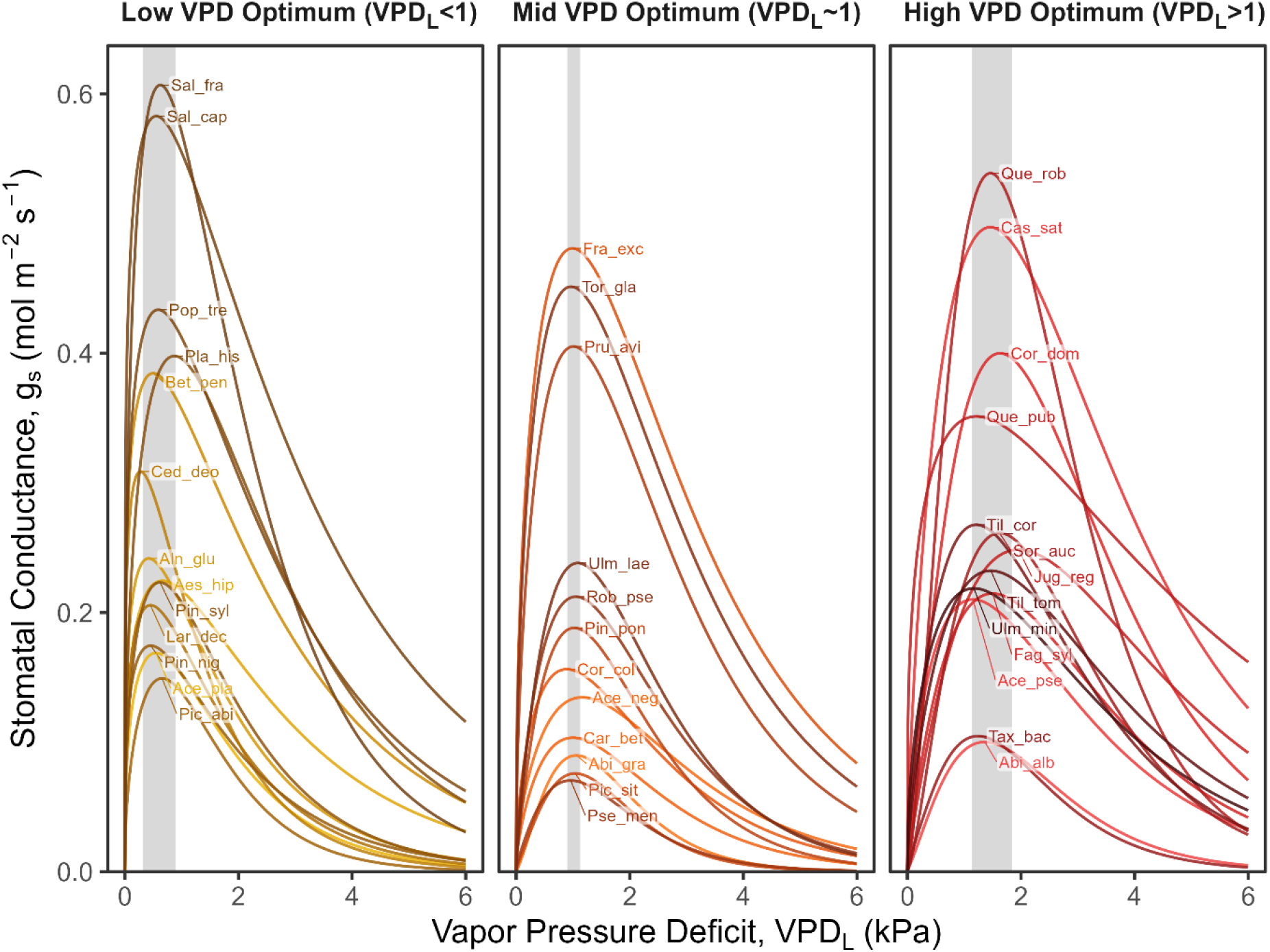
Species-specific responses of stomatal conductance (g_s_) to leaf-to-air vapor pressure deficit (VPD_L_). Curves represent expected population-level mean responses calculated via posterior predictions from the Bayesian model, holding ambient light (Q_amb_) constant at the mean. Panels partition species into three equal-sized groups (tertiles) based on the VPD optimum (VPD_opt_), the VPD_L_ value at which maximum stomatal conductance is predicted. Shaded grey backgrounds indicate the range of species’ VPD_opt_ values. Line colours are used for visual distinction. Table 1 gives species abbreviations.

Species differed in the VPD_L_ at which maximum conductance was reached, the magnitude of this maximum, and the decline in g_s_ with atmospheric demand. To quantify these differences, we derived four stomatal behaviour metrics that together capture the key facets of the response curves (see Figure 1c, Table 2 for their definition). Figure 3 summarises these metrics across all species considered here (see Table S6 for species-specific values), revealing four descriptive regions of stomatal behaviour space: high-capacity early-peaking, high-capacity late-peaking, low-capacity early-peaking, and low-capacity late-peaking species. Conifer species were predominantly located in the low-capacity early-peaking quadrant, whereas broadleaved species spanned all four behavioural types (Figure 3). Substantial interspecific variation was present within each response type, highlighting that stomatal behaviour varied along continuous gradients rather than forming distinct clusters. g_max_ varied ca. eightfold across species (0.077 - 0.648 mol m^-^² s^-1^; Figure S7), with conifers showing a statistically clear lower average g_max_ at half of that of broadleaved species (0.16 vs. 0.34 mol m^-^² s^-1^; Figure S8). Nevertheless, some conifers, such as *Cedrus deodara* reached values comparable to those of many broadleaved species, and some broadleaved species such as *Carpinus betulus* or *Acer negundo* showed g_max_ on or below the level of conifer species.

**Figure 3:**
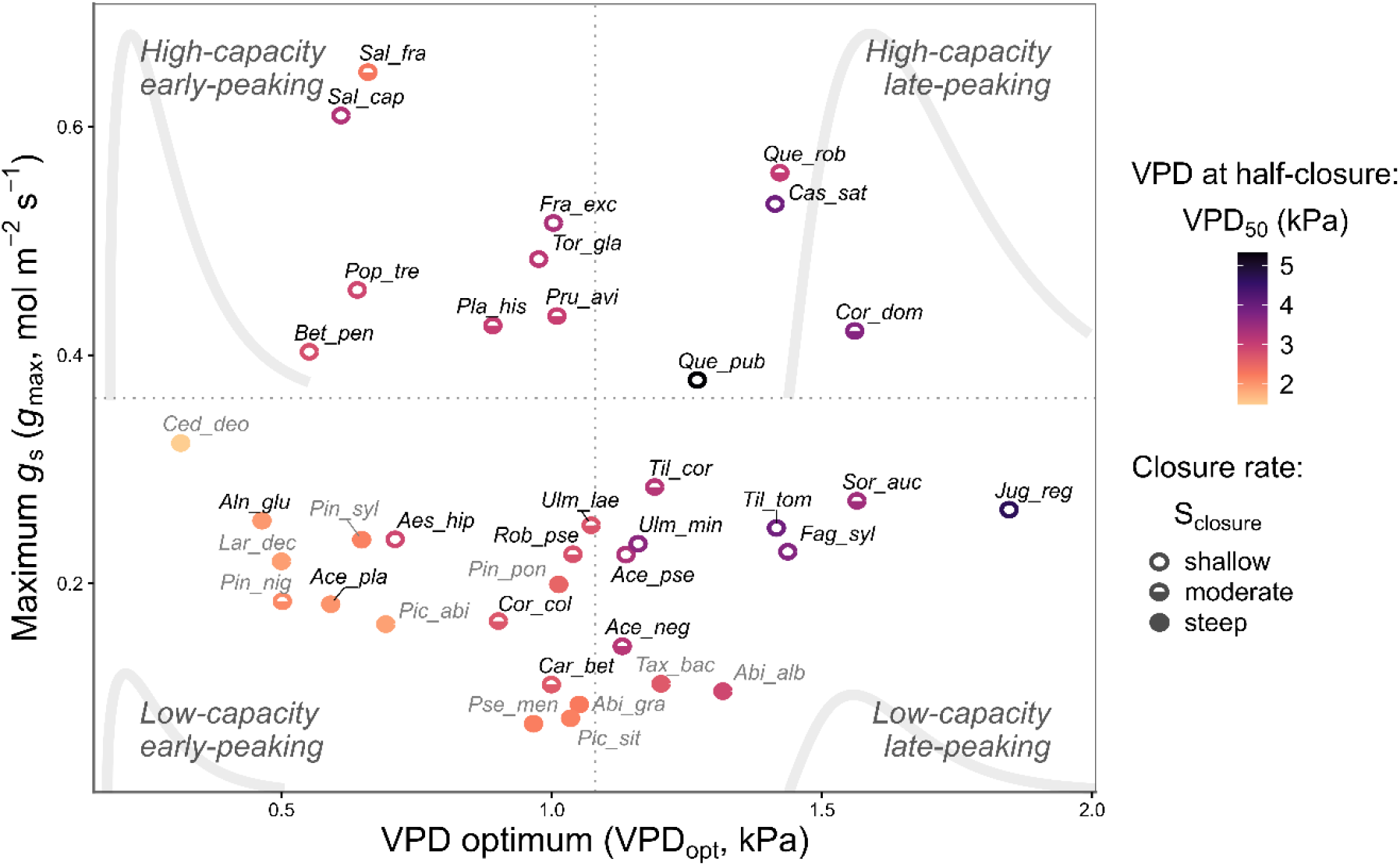
Species-specific stomatal behaviour metric estimates: maximum stomatal conductance (g_max_), VPD optimum (VPD_opt_), VPD at half-closure (VPD_50_) and stomatal closure rate (S_closure_). Points represent posterior means for each species. g_max_ and VPD_opt_ are shown on the y- and x-axis, respectively, while VPD_50_ is shown in the colour gradient and the information on S_closure_ is summarised in three point shape groups. Species labels are shown in grey for conifers and in black for broadleaved species. Grey dotted lines indicate the median values of g_max_ and VPD_opt_, dividing the space into four response types: high-capacity early peaking, low-capacity early-peaking, high-capacity late-peaking, and low-capacity late-peaking. The typical shapes of these response curves are shown in light grey in their respective space. See Table 1 for species abbreviations, and Table S6 for exact values of the metrics.

The VPD_opt_ ranged from 0.31 to 1.85 kPa, whereas the VPD_50_ ranged from 1.48 to 5.32 kPa (Figure S7), indicating substantial interspecific variation in the onset and progression of stomatal closure. Species that reached their peak g_s_ at higher VPD_L_ (high VPD_opt_), generally also dropped their conductance to 50% of their g_max_ at a higher VPD_L_ (high VPD_50_) (Pearson *r* = 0.74, Figure S9), suggesting a coherent VPD response syndrome. However, some species like *Salix caprea* both reached their g_max_ at a very low VPD but were subsequently very slow to close their stomata with rising VPD (Figure 3). VPD_50_ was consistently lower in conifers than in broadleaved species, with a difference of almost 1 kPa between clades (Figure S8), whereas the difference in VPD_opt_ was smaller (0.22 kPa) and not statistically clear (Figure S8). The decline slope of the curve (S_closure_), a proxy for the closure rate with rising VPD_L_, also varied substantially between species (range: very fast reacting species 0.55 mol m^−2^ s^−1^ kPa^−1^ vs. slow reacting species 0.17 mol m^−2^ s^−1^ kPa^−1^; Figure S7). Conifers generally had a steeper closure rate than broadleaved species (Δ=0.16 mol m^-2^ s^-1^ kPa^-1^, Figure S8).

Species that showed a higher VPD at half-closure also generally had a less steep S_closure_ (Pearson *r* = - 0.85; Figure S9, S10). Species with lower g_max_ showed a tendency to close their stomata more rapidly (Pearson *r* = -0.51; Figure S9). Even though low-capacity early-peaking species generally showed a steep stomatal closure with increasing VPD, this pattern was not universal; for instance, *Aesculus hippocastanum* combined a relatively low g_max_ with a comparatively shallow closure rate (Figure 3). VPD_opt_ and VPD_50_ were strongly correlated (Pearson *r* = 0.74; Figure S9), suggesting that these metrics capture closely related and partly redundant aspects of the VPD response. In contrast, g_max_ showed no relationship with VPD_opt_ and only a weak relationship with VPD_50_ (Pearson r= -0.31; Figure S9), suggesting maximum gas exchange rates and VPD sensitivity response metrics represent distinct dimensions of stomatal behaviour. This was further supported by the PCA (Figure S10; PC1 = 62.4%, PC2 = 27.7% of variance explained), in which VPD_opt_, VPD_50_, and S_closure_ loaded primarily on PC1 (0.45–0.61), whereas g_max_ loaded only weakly on PC1 (0.31) but dominated PC2 (0.76), consistent with g_max_ representing an independent dimension from the VPD sensitivity metrics.

### Physiological drought responses reflect stomatal sensitivity to VPDL

The VPD optimum and VPD at half-closure were associated with drought-induced differences in leaf δ¹³C. Species with a higher VPD optimum showed a smaller shift in leaf δ¹³C between the extreme drought year 2018 and the wet reference year 2021, indicating stronger physiological drought responses (Figure 4b).

**Figure 4:**
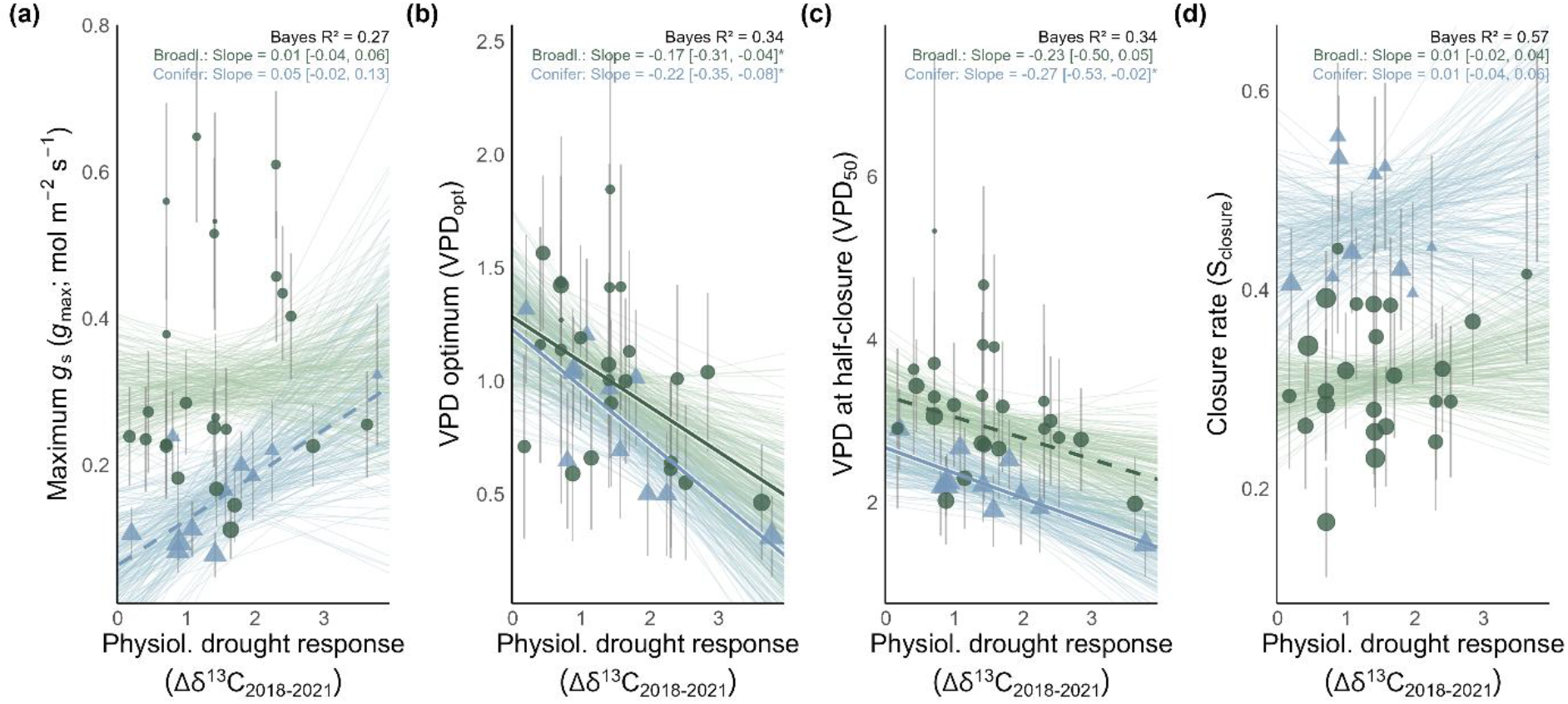
Relationship between stomatal behaviour metrics and physiological drought response (leaf Δδ^13^C_2018-2021_, difference between drought year 2018 and reference year 2021). Bold lines indicate statistically clear (95 % credible interval) and bold dashed lines marginally statistically clear (66% credible interval) effects; the posterior uncertainty cloud is based on 200 random draws from the model parameters. Error bars indicate ± CI around individual species means; point size is scaled inversely to variance (1/SD) to reflect parameter precision in model weighting. Please note that R² values reflect total model fit, which in some cases is driven primarily by intercept differences between tree groups (conifer/broadleaved) rather than by slope relationships with the main predictor. See Table S9 for detailed model summaries.

This relationship was moderately strong (R² = 0.34) but notably consistent across broadleaved and conifer species (Figure 4b). A similar pattern was found for the metric of VPD at half-closure, VPD_50_, with more tolerant species (higher VPD_50_) having shown less physiological drought responses in 2018, though the relationship was only statistically clear for conifers (Figure 4c). Neither g_max_ nor S_closure_ showed a statistically clear relationship with leaf Δδ^13^C, although g_max_ showed a tendency toward a positive relationship for conifers (Figure 4a,d). Overall, species that can tolerate higher VPD_L_ and function optimally under these conditions also maintain a lower carbon isotope discrimination during a hot drought, a pattern that held consistently across both broadleaved species and conifers.

### Leaf functional traits associated with stomatal behaviour

Testing the relationship of the four stomatal behaviour metrics and seven leaf traits revealed that whilst some leaf traits could predict maximum g_s_ for broadleaved species, none could reliably predict VPD optimum, VPD at half-closure, or the closure rate (Figure 5).

**Figure 5:**
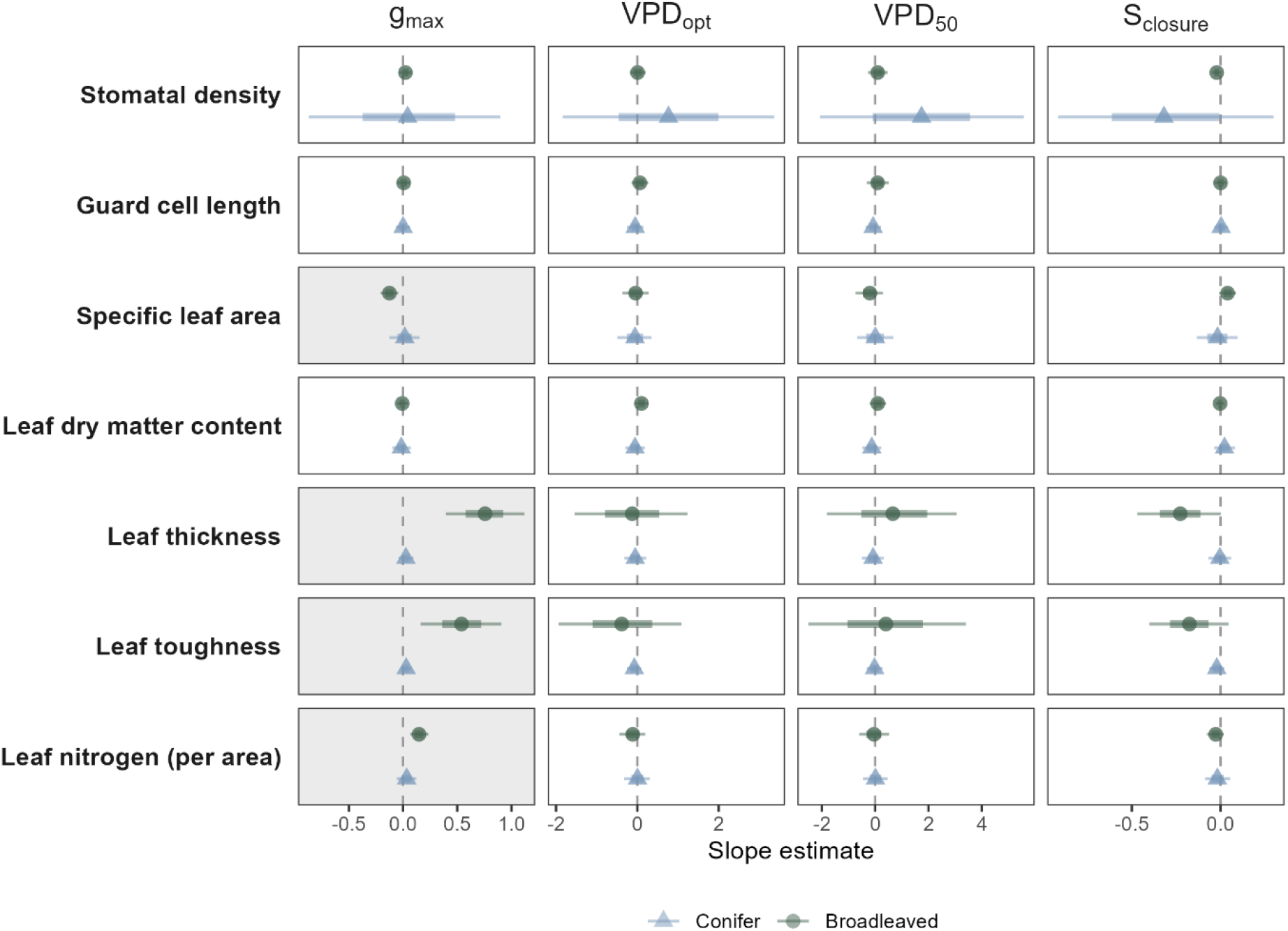
Standardized slope estimates from Bayesian linear models relating leaf functional traits to stomatal behavioural metrics across broadleaved and conifer species. For each metric (maximum conductance (g_max_), VPD optimum (VPD_opt_), VPD at half-closure (VPD_50_) and stomatal closure rate (S_closure_)), we fitted a separate model for each scaled trait (z-scored across all species) in interaction with the tree group (broadleaved/conifer) as predictor, allowing both slopes and intercepts to differ between groups. Each panel shows the posterior median slope and 95% credible interval for broadleaved (circles) and conifer (triangles) species separately, with the 66% credible interval shown as a thicker point range. Positive slopes indicate that species with higher trait values tend to show higher values of the corresponding stomatal behavioural metric; negative slopes indicate the inverse. Grey shading indicates panels where at least one group’s 95% credible interval excludes zero (statistically clear effect). See Table S9 for model summaries of all 28 models.

Neither stomata density nor guard cell length showed a statistically clear effect on the g_max_. Lower SLA was associated with higher g_max_ (Bayes R² = 0.43; broadleaved slope = -0.13 [-0.21, -0.04]), contrary to our expectations. Species with thicker leaves (Bayes R² = 0.51; broadleaved slope = 0.75 [0.39;1.16]) and tougher leaves (Bayes R² = 0.38; broadleaved slope 0.54 [0.14, 0.94]) also showed a higher g_max_, as did species with a higher leaf nitrogen content (Bayes R² = 0.51; broadleaved slope = 0.15 [0.06, 0.24]), with all effects statistically clear for broadleaved species only (Figure 6).

**Figure 6:**
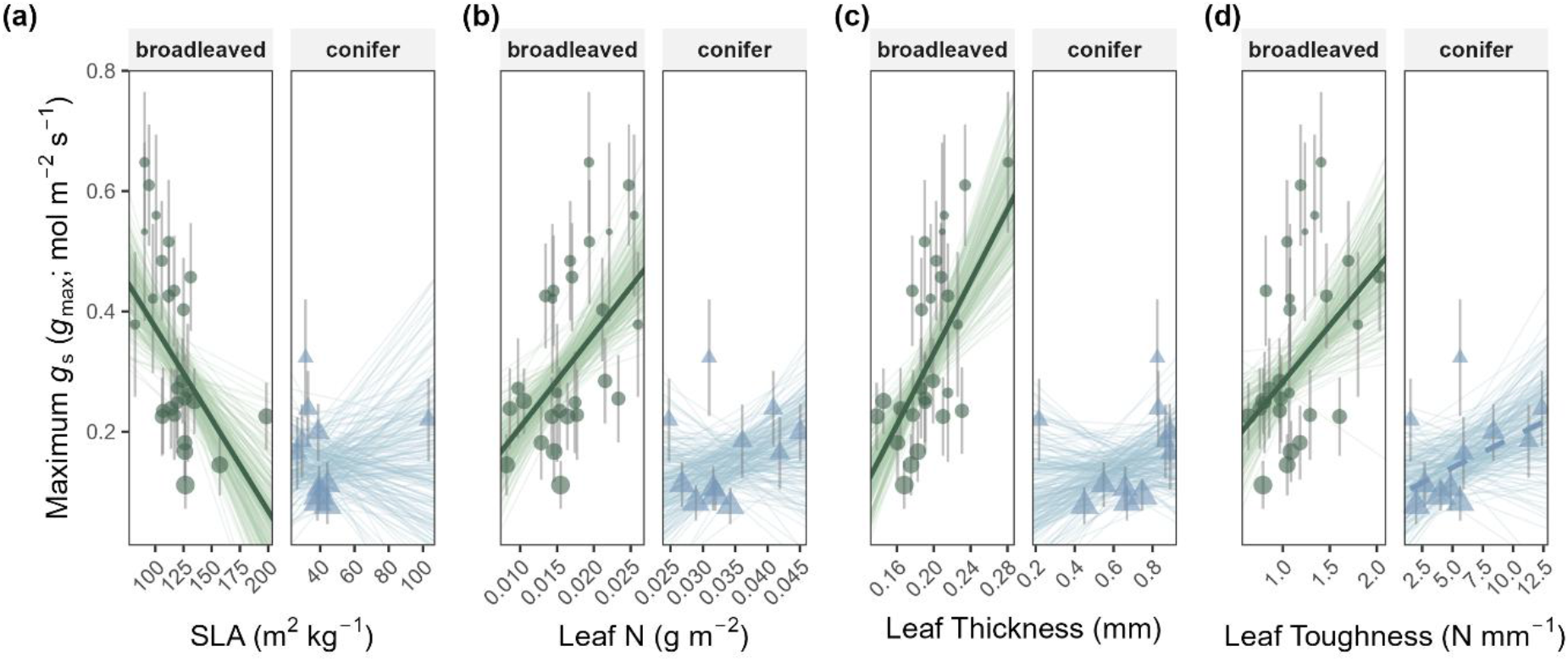
Relationship between maximum stomatal conductance (g_max)_ and selected leaf functional traits. Models included an interaction between tree group and leaf trait; broadleaved species and conifers are shown in separate panels because of their different trait ranges. Panels show (a) specific leaf area (SLA), (b) leaf nitrogen per area, (c) leaf thickness, and (d) leaf toughness. Larix decidua, the only deciduous conifer in our species selection, was an outlier in some models due to its exceptionally high SLA (103.3 m^2^ kg^-1^) and very thin needles (0.22 mm). Bold solid lines indicate population-level mean trends with 95% credible intervals excluding zero (bold dashed lines for 66% credible intervals). the posterior uncertainty cloud is based on 200 random draws from the model parameters. Error bars indicate ± CI around individual species means; point size is scaled inversely to variance (1/SD) to reflect parameter precision in model weighting. Bayes R^2^: (a) 0.43, (b) 0.45, (c) 0.51, and (d) 0.38. Please note that R² values reflect total model fit, which in some cases is driven primarily by intercept differences between tree groups (conifer/broadleaved) rather than by slope relationships with the main predictor. See Table S9 for model summaries.

Neither stomatal density nor guard cell length could predict VPD_opt_, VPD_50_, or S_closure_ (Figure 5). However, some tendencies emerged: species with higher stomatal density tended toward higher VPD_opt_, and VPD_50_ (Figure S11a,d). Stomatal density tended to be negatively associated with closure rate, such that species with a higher density of stomata closed more slowly with rising VPD, in both broadleaved and conifer species (Figure S11g).

In contrast to their ability to explain maximum g_s_, neither SLA and leaf nitrogen, nor leaf thickness and toughness could predict any of the stomatal sensitivity metrics to VPD_L_. Note, however, that among broadleaved species, (i) higher SLA species tended to also display lower VPD at half-closure (Figure S11e), and faster stomatal closure with rising VPD_L_ (Figure S11h); and (ii) thicker-leaved species tended to close their stomata more slowly with rising VPD_L_ (66% credible interval; Figure S11i). Because these associations were not statistically clear, our results overall suggest that stomatal sensitivity to VPD_L_ is largely independent of the leaf functional traits examined here, across both broadleaved species and conifers.

## Discussion

Based on an extensive collection of stomatal conductance data for 38 temperate tree species in the field and under the same conditions, we found substantial interspecific variation in stomatal behaviour in response to VPD_L_. Species differences in VPD sensitivity were associated with drought-induced changes in leaf Δδ^13^C, linking stomatal behaviour to past physiological drought responses. While some leaf anatomical and economic traits explained variation in maximum stomatal conductance, the stomata’s response to VPD_L_ was largely decoupled from commonly measured leaf traits.

### Metrics describing stomatal behaviour

Compared with previous approaches that characterized observed VPD-induced stomatal downregulation from a reference point (e.g., g_max_ or the reference g_s_ at a set VPD) through a single sensitivity parameter (e.g., Oren et al., 1999), we propose to resolve stomatal behaviour into four physiologically distinct dimensions: *how much* gas exchange a species is capable of (g_max_), *when* atmospheric demand begins to constrain it (VPD_opt_), *how long* substantial gas exchange can be maintained (VPD_50_) and *how abruptly* the stomata close (S_closure_).

The substantial interspecific variation in the VPD optimum found in our study (Figure 2, Figure 3) highlights that the choice of the reference conductance (g_s,ref_) from which to assess stomatal downregulation in response to increasing VPD is critical. Normalising species’ response to a common VPD (e.g., 1 kPa; Oren et al. 1999) can place g_s,ref_ at different positions along species-specific response curves, depending on their VPD_opt_. Using a species-specific reference point, as in our approach, may therefore provide a more meaningful basis for comparing stomatal sensitivity to VPD across species.

The four dimensions of stomatal behaviour appear to represent two independent axes. g_max_ primarily reflects the capacity for gas exchange and thus the acquisitive aspect of plant functioning, whereas VPD_opt_, VPD_50_, and S_closure_ rather capture stomatal sensitivity to water loss. g_max_ was largely unrelated to the VPD_L_ thresholds of stomatal closure (VPD_opt_, VPD_50_) and only marginally related to the closure rate (S_closure_; Figure S9, S10). We therefore found no evidence of the classical safety-efficiency trade-off, according to which species with higher g_max_ ought to be more prone to closure (Henry et al., 2019). Instead, stomatal sensitivity to VPD_L_ appears to form a distinct syndrome that can vary relatively independently from stomatal capacity (Figure S10), a pattern consistent with previous trait-based analyses of stomatal control in subtropical trees (Kröber & Bruelheide, 2014; Schnabel et al., 2024).

In our study, the sensitivity metrics were not independent of each other, pointing to coordinated control strategies. For instance, the negative correlation between VPD_50_ and S_closure_ is plausible, as risk-averse species may close their stomata early and abruptly, whereas risk-tolerant species may pursue gas exchange far into dry conditions and close gradually. This relationship is also partly inherent to the metrics’ definitions, because both pertain to a decline in conductance relative to *g*_max_. Similarly, the positive correlation between VPD_opt_ and VPD_50_ illustrates how the same capacity to maintain stomatal opening as the atmosphere dries underlies both metrics. Nevertheless, deviations from the aforementioned relationships demonstrate nuance in the aspects captured by these sensitivity metrics, supporting their complementary use. For example, the low VPD_opt_ but relatively high VPD_50_ of *Salix caprea* may reflect the opportunism of a pioneer species that maintains high carbon gain during favourable periods, via relatively less reactive stomata in its typically mesic or moist habitats (Enescu et al., 2016).

Our multidimensional framework also revealed clear differences between conifers and broadleaved species (Figure S8). As expected, conifers showed lower g_max_ than broadleaved species, consistent with their conservative leaf economics (Reich, 2014; Wright et al., 2004) and anatomical constraints on water transport, including narrow tracheid-based xylem and simple needle venations, which limit hydraulic conductance and thus maximum gas exchange (Brodribb et al., 2007; Sperry et al., 2006). Conifers often also have sunken, wax-plugged stomata, which adds resistance to water loss (Šantrůček, 2022). Although gymnosperms possess the biochemical machinery for ABA signalling, short-term stomatal responses to increasing VPD are thought to be governed predominantly by passive hydraulic mechanisms (Brodribb & McAdam, 2011, 2017; Franks & Farquhar, 2007), potentially explaining their rapid stomatal closure once a critical leaf water threshold is crossed (Figure S8, McAdam & Brodribb, 2014, 2015). However, because conifers and broadleaved species were measured with different porometer devices (LI-600N vs. LI-600), differences in boundary layer conductance estimation may also have contributed to the observed g_max_ differences.

Beyond broad distinctions between conifers and broadleaved species, their positions in the multidimensional trait space reflected their ecological strategies. Pioneer species (e.g. *Salix caprea*, *Populus tremula*, or *Betula pendula)* generally combined high g_max_ with a low VPD_opt_. Likewise, species associated with riparian or consistently moist habitats tended to have low VPD_opt_ (e.g. *Alnus glutinosa, Salix fragilis, or Acer platanoides*), which may reflect weaker constraints on stomatal opening in species adapted to more reliable water supply. In contrast, drought-tolerant or sub-Mediterranean species (e.g. *Juglans regia, Tilia tomentosa, Castanea sativa*, or *Quercus pubescens)* generally had higher VPD_opt_. *Fagus sylvatica* also showed a very high VPD_opt_ and high VPD_50_, consistent with its reported anisohydric tendencies (Bréda et al., 2006; Jonard et al., 2011; Köcher et al., 2009; Leuschner et al., 2022; Niemczyk et al., 2024), which may become particularly consequential during hot drought, when high VPD coincides with temperatures that can impair hydraulic and photosynthetic functioning (Bär et al., 2018; Geßler et al., 2007; Leuschner, 2020).

These ecological interpretations should be considered in light of the environmental conditions under which our metrics were derived. Our metrics were based on leaf-to-air VPD (VPD_L_) under non-limiting soil moisture conditions and therefore primarily describe species responses to atmospheric demand. We used VPD_L_ because it more accurately reflects the leaf boundary micrometeorology than ambient VPD, given that leaf temperature deviates from air temperature due to radiative heating and transpirational cooling (Jones, 2013; Lian et al., 2026; Monteith & Unsworth, 2007). The deviation from site- to leaf-level conditions was substantial in our dataset: while ambient VPD did not exceed 3.02 kPa, values of VPD_L_ of up to 6.11 kPa were recorded (Figure S12). Leaves were indeed often warmer than the surrounding air, owing to strong radiative heating. Nevertheless, species maintaining higher stomatal conductance, like *Quercus pubescencs,* also might have cooled their leaves more strongly and experienced lower VPD_L_ than species more sensitive to the same atmospheric conditions (Figure S12).

### Stomatal sensitivity reflects integrated physiological drought responses

Our results showed that metrics of stomatal sensitivity, namely the VPD optimum and the VPD at half-closure, are linked to physiological drought responses in a hot drought year. Species with a lower VPD optimum (earlier stomatal downregulation as atmospheric demand increased), showed greater drought-induced increases in leaf Δδ¹³C, consistent with stronger stomatal limitation of CO₂ uptake and reduced intercellular CO₂ concentrations during drought. We interpret this finding as an independent validation of our stomatal behaviour metrics, whose association with drought-induced changes in leaf Δδ^13^C confirms that they capture ecologically relevant differences among species. We note that Δδ^13^C is not a clear-cut readout of stomatal closure: intercellular CO_2_ concentrations also depend on photosynthetic capacity, and the elevated temperatures characteristic of hot droughts can independently constrain photosynthetic activity, potentially contributing to the observed δ^13^C signal (Cernusak et al., 2013; Scafaro et al., 2023). The strength of the relationship may partly reflect the exceptional severity of the 2018 drought across Central Europe and at our site (Figure S6; Bastos et al., 2020; Rakovec et al., 2022; Zscheischler & Fischer, 2020). Thus, although leaf Δδ^13^C integrates multiple physiological processes over the growing season, its association with a within-day, high-resolution measure of species-specific stomatal behaviour strategy implies that stomatal sensitivity to VPD is indeed a genuine and detectable contributor to physiological drought responses.

An open question is whether our metrics also reflect stomatal regulation in response to soil drying specifically, since water-use strategies that appear conservative at a daily timescale do not necessarily remain conservative over the longer timescale relevant to soil drying (Koehler et al., 2023). The severe 2018 drought was a compound event with both high VPD and reduced soil moisture (Bastos et al., 2020; Zscheischler & Fischer, 2020) and therefore the relationship with Δδ^13^C reflects integrated responses to both stressors. While our results indicate that species differences in VPD_L_ responses contribute to species drought behaviour, their relative importance compared with soil drought responses cannot be resolved here. Experiments are starting to disentangle soil water stress from VPD effects in the field (e.g., *VPDrought,* https://vpdrought.wsl.ch), but synthesizing their relative importance across a broad set of species will remain challenging.

Beyond the leaf level, physiological drought responses rarely stay an isolated signal but typically propagate into the whole-tree carbon economy, since elevated temperatures and stomatal closure limit photosynthetic carbon uptake, and this reduction in assimilation is commonly reflected in reduced growth during and after drought years (Jucker et al., 2017; Kretz et al., 2025; Schnabel et al., 2022). Our stomatal behaviour metrics therefore capture more than a snapshot of daily gas exchange regulation, rather reflecting species-level strategies with potential consequences for carbon assimilation and growth during drought.

### Leaf traits of the fast-slow continuum can explain variation in gmax

In contrast to our expectations, g_max_ was not correlated with stomatal density or guard cell length, despite diffusivity principles clearly linking total pore area to leaf gas-exchange capacity (Farquhar & Sharkey, 1982; Franks & Beerling, 2009; Franks & Farquhar, 2001). Stomatal density and guard cell length were negatively correlated across species (Figure S13), in line with the commonly observed stomatal size-density trade-off (de Boer et al., 2016; Liu et al., 2023). The relationship between stomatal anatomy and realized gas exchange is more complex in empirical studies than predicted from mathematical and physical considerations. Although anatomical estimates and measured operational maximum conductance appear positively related across a broad range of species, only a fraction of the theoretical anatomical value is typically realized (McElwain et al., 2016; Murray et al., 2020); this realised fraction appears to be both context-dependent and notably small at current atmospheric CO_2_ concentrations (Dow et al., 2014). This hints at g_max_ being heavily governed by physiological control in addition to anatomical constraints, such that variation in stomatal traits does not necessarily translate into differences in field-measured g_max_.

Variation in g_max_ across our study species was, at least for broadleaved species, rather explained by traits reflecting leaf economy (Wright et al., 2004), specifically leaf nitrogen, specific leaf area, leaf thickness, and leaf toughness (Figure 6). Based on the fast-slow continuum, we initially expected thinner leaves with higher SLA, associated with a fast-growth resource-acquisitive strategy (Reich, 2014) to support higher g_max_. We found the opposite: low-SLA, thick, and tough leaf tissue correlated with higher g_max_. This pattern makes sense when considering the area-based expression of g_max_, because thicker leaves contain a greater volume of photosynthetic tissue stacked beneath each unit of leaf surface. Greater structural packing results in higher nitrogen per unit area, reflecting investment in the photosynthetic machinery that may elevate internal CO_2_ demand and thereby require higher g_max_ per unit volume to keep the Calvin cycle adequately supplied (Evans, 1989; Schulze et al., 1994). Further, leaves with high LDMC, greater thickness, and higher toughness represent a greater investment in leaf construction (Niinemets, 1999). We speculate that such structural investment is linked to increased leaf hydraulic conductance, as higher vein density and greater xylem investment enhance the delivery of water to evaporating sites and sustain the water flux required for high stomatal opening (Brodribb et al., 2007; Nardini, 2022; Sack & Holbrook, 2006; Sack & Scoffoni, 2013). In this context, low-SLA, thick and tough leaves do not necessarily represent a conservative ‘slow’ strategy; rather, they may achieve high gas exchange per unit area by concentrating both photosynthetic and hydraulic investments within a smaller leaf area. For conifers, we observed similar but much weaker directional trends (Figure 6). These likely reflect the limited sample size (11 conifer species), the influence of *Larix decidua*, the only deciduous conifer with a distinct physiology (Matyssek, 1986), and constrained trait variation in conifers, which generally have low SLA and long-lived, structurally reinforced needles (Reich, 2014). In addition, for some conifer species, curve-derived g_max_ occurred at very low VPD_L_, not captured in our measurements (Figure S5), indicating that *g*_max_ could be underestimated.

### Stomatal VPDL sensitivity is decoupled from anatomical and economic leaf traits

Although leaf traits appear to constrain g_max_, they provide little insight into how stomata regulate gas exchange in response to VPD_L_. None of them explained the VPD optimum, VPD at half-closure, or stomatal closure rate, suggesting that stomatal sensitivity to VPD_L_ is largely independent of stomatal anatomy and leaf economy, at least based on the traits considered here.

We expected species with fewer and larger stomata to experience higher water loss when evaporative demand is high (Drake et al., 2013; Kröber & Bruelheide, 2014; Wang et al., 2015), and hypothesised that these species would close their stomata at lower VPD_L_, but found no evidence for this. The weak tendency towards steadier closure in species with higher stomatal density may instead indicate that numerous small stomata enable more gradual and spatially heterogeneous regulation, for example through asynchronous closure across the leaf surface, e.g. beginning in less favourable regions such as areas further away from major veins (Lawson and Blatt 2014; Mott and Buckley 1998). In contrast, species with fewer, larger stomata may lack this capacity for incremental closure and transition more abruptly towards closure.

Similarly, we found only weak evidence that broadleaved species with a higher SLA have a lower VPD at half-closure and a higher closure rate, as expected by the LES framework (Reich, 2014). This expectation is derived from the assumption that acquisitive, fast-growing species typically prioritise rapid resource turnover and close their stomata when met with declining water supply, thereby protecting their hydraulic system from tension-induced failure (Henry et al., 2019). However, our data do not support the notion that leaf economic traits generally predict stomatal sensitivity. Similarly, previous trait-based studies found no association between leaf economic traits and stomatal sensitivity, with stomatal control gradients appearing orthogonal to the fast-slow leaf economics spectrum (Kröber & Bruelheide, 2014; Sachsenmaier et al., 2025; Schnabel et al., 2021, 2024). Further, we found no relationship between leaf thickness and VPD optimum or half-closure threshold, despite expectations that thicker leaves might buffer water status through greater hydraulic capacitance (Blackman & Brodribb, 2011). However, the relationship between hydraulic capacitance and stomatal sensitivity may depend on species-specific hydraulic thresholds (Martinetti et al., 2026), potentially obscuring relationships across diverse species pools. The weak negative relationship between thickness and closure rate observed here may nevertheless suggest that greater water storage capacity reduces the need for rapid stomatal closure (Fu et al., 2019; Qie et al., 2024).

The lack of relationships between VPD sensitivity metrics and commonly measured leaf traits highlights that stomatal behaviour likely depends on traits more directly linked to plant water regulation or leaf gas exchange. The carbon gain from maintaining stomatal opening under high VPD may depend on how well photosynthesis is maintained at the high temperatures that often accompany high VPD, as reflected by traits such as the optimum temperature for photosynthesis (Crous et al., 2022). Relevant hydraulic trait candidates include leaf turgor loss point, maximum leaf hydraulic conductance, leaf capacitance and vein density (Bartlett et al., 2012; Blackman et al., 2018; Brodribb & Holbrook, 2003; Sack & Holbrook, 2006). Moreover, stomatal regulation is increasingly understood as coordinated with whole-plant hydraulics, including roots and stems, rather than being governed on the leaf level in isolation (Hartmann et al., 2021). How long a species keeps its stomata open under rising VPD is unlikely to be completely independent of the risk this poses to the xylem through embolism (Martin-StPaul et al., 2017), linking stomatal behaviour to a species’ embolism resistance, hydraulic safety margin and whole-plant hydraulic capacitance. Such hydraulic traits have been shown to coordinate with regulation of leaf water potential along the isohydric-to-anisohydric spectrum (Martínez-Vilalta & Garcia-Forner, 2017), further mediated biochemically through species differences in foliar abscisic acid (ABA) dynamics and sensitivity (Brodribb & McAdam, 2013; Sussmilch et al., 2017). However, the extent and consistency of this coordination across species remain unclear, because a comparable description of stomatal behaviour across diverse species has so far been lacking. Future work at the same site aims therefore to test how stomatal behaviour metrics are coordinated with hydraulic traits across the whole tree.

Taken together, our comparison of 38 tree species demonstrates substantial interspecific variation in stomatal behaviour that is relevant to understanding species’ drought responses, but cannot be predicted by commonly measured anatomical and economic leaf traits. The large variation revealed here highlights a challenge for models that represent stomatal responses using fixed or generalised sensitivity responses. Our results further show that stomatal behaviour is multidimensional. Absolute gas-exchange capacity and stomatal regulation under increasing atmospheric demand represent distinct behavioural dimensions that should be explored systematically and separately, to make for a more comprehensive representation of how tree species balance carbon gain against water loss. Our stomatal behaviour metrics complement existing trait-based approaches by capturing dynamic aspects of stomatal drought responses. In particular, the VPD optimum emerges as a promising indicator to help forecast species’ future drought stress. Finally, the thorough characterisation of stomatal behaviour may help reduce uncertainty in vegetation and land-surface models, where stomatal regulation remains simplified (Grossiord et al., 2020; Medlyn et al., 2011; Sabot et al., 2022) and improve predictions of forest responses to increasing atmospheric demand.

## Acknowledgements

We thank the members of the working group Systematic Botany and Functional Biodiversity (Leipzig University) for their exceptional support during the demanding field campaign, especially the technical staff, Julia van Braak, Tom Künne and Nicole Nabel, and the student helpers Jonas Pohlmann, Greta Zahner, Melanie Hanfstängel, Jessica Esser and Carolin Hensel. We thank Julia Olbrich and Greta Zahner for their thesis work on leaf carbon isotope signals, and Heiko Moossen and Heike Geilmann at the BGC-IsoLab at MPI Jena for the isotope measurements. We thank Jonas Franke and Benjamin Rosenbaum for discussions on statistics. This study was supported by the International Research Training Group TreeDì jointly funded by the Deutsche Forschungsgemeinschaft (DFG, German Research Foundation) – 319936945/GRK2324 and the University of Chinese Academy of Science (UCAS). Open Access funding enabled and organised by Projekt DEAL.

## Author contributions

LS, RR and CW conceived the ideas and designed the study. LS, CA, RR, LK, AK and FS collected the data. CA performed preliminary analyses. LS cleaned and analysed the data, supported by RR, DSC, IS and MS. MS provided scientific guidance throughout the study. CW supervised the project and provided funding. LS wrote the first draft of the manuscript. All authors contributed critically to the manuscript and approved the final version.

## Data availability

The data supporting the findings of this study are available in Zenodo. The stomatal conductance data (and associated site climate data) are available at https://doi.org/10.5281/zenodo.22308419. Stomatal anatomy trait data of gymnosperms are available from Sachsenmaier et al. (2026b) (https://doi.org/10.5281/zenodo.21098915) and corresponding data for angiosperms are available from Kretz, Sachsenmaier et al. (2026) (https://doi.org/10.5281/zenodo.22117766). Leaf carbon isotope data are available from Kahl, Schnabel et al. (2026) (https://doi.org/10.5281/zenodo.22284412), and leaf economics trait data are available from Kretz et al. (2026) (https://doi.org/10.5281/zenodo.21821305). Some datasets remain restricted during peer review; private access links have been provided to the editorial office for reviewers and will be made public upon publication. The R code supporting the analyses is available on GitHub: [https://github.com/LenaSachsenmaier/Stomatal_Behaviour_Analysis/releases/tag/v1.0.0].

