## Supplementary Notes, Tables and Figures for "Stomatal sensitivity to VPD across 38 tree species reflects past drought responses but is little explained by stomatal and leaf economic traits"

The following Supporting Information is available for this article:

### Supplementary Methods Section

#### Note S1: Adjustments for measurements of needle-leaved species

For gymnosperms, needle widths were preconfigured in the instrument software based on species-specific averages measured in advance (Table S2). Because the needles of several species were too short to span the entire length of the measurement chamber, we mathematically adjusted the leaf area post-measurement, using species-specific correction factors derived from preliminary tests, which quantified the average percentage of the chamber length occupied by the needle (Table S2). Custom gaskets were used to match varying needle diameters and ensure chamber sealing.

#### Note S2: Quality control of $g_s$ values

Environmental factors were verified to be within valid ranges of the devices' readings ($VPD_L \geq 0.05$ kPa, relative humidity ≤ 95%), and measurements with excessive chamber leakage were excluded (>10% for the LI-600, >15% for the LI-600N; the higher threshold for the needle porometer reflects the difficulty of achieving low leakage in a closed chamber with a needle insertion). Physiologically implausible gas exchange values (i.e., negative $g_s$) were set to zero only when they coincided with very low leakage (<5%) or with negative fluorescence and electron transport readings indicating negligible conductance (n=50); all other implausible values were removed (n=21). Only leaves with at least six reliable measurements over the course of the day were retained, resulting in 4-7 leaves per tree and 18-21 leaves per species. Each tree contributed 20-70 measurements throughout the day, resulting in 111–188 measurements per species and a total of 6002 observations.

#### Note S3: Stomata density and size measurements

Stomatal imprints were prepared using transparent nail varnish. Conifer needles were pre-treated with hot water and detergent to remove epicuticular waxes obscuring stomata, following the protocol of Sachsenmaier et al. (2026). Imprints were taken from the abaxial surface of angiosperm leaves and across the relevant needle surfaces of conifers to match the respective gas exchange measurements. For angiosperms, stomata were counted in three randomly selected fields of view per imprint at 400x magnification using a light microscope (DM300, Leica). Conifer stomata were counted at 100x magnification following Sachsenmaier et al. (2026) using digital images analysed in ImageJ. Stomatal density, expressed as number of stomata per $mm^2$, represents the abaxial surface for angiosperms and total projected needle surface for conifers.

Guard cell length (GCL) was measured at 1000x magnification. For each imprint, one centrally located stoma from three randomly selected fields of view was measured, either directly using a calibrated ocular micrometer (angiosperms) or digitally using ImageJ (conifers). All measurements were performed by a single observer (separate for angiosperms and conifers).

### Supplementary Tables

*Table S 1: Environmental conditions during the measurement days.*
*For each of the eight measurement dates, the time of the first and last measurement (start time, end time) are given in Central European Time (CET) and the number of tree individuals measured during this day (in sum: n=114) is indicated. For each day-specific measurement window, median values are reported for air vapour pressure deficit (VPD; kPa), air temperature (°C), relative humidity (%), wind speed (m $s^{-1}$), global radiation (W $m^{-2}$], diffuse radiation fraction relative to global radiation (diffuse/global), and soil volumetric water content (%) at 15 cm and 50 cm depth. Data were obtained from an on-site climate station equipped with a WXT520 weather sensor (Vaisala, Finland), a SPN1 sunshine pyranometer (Delta-T Devices Ltd, England) and ML3 ThetaProbe soil moisture sensors (Delta-T Devices, UK), recording at 10-minute intervals. Soil water content was measured using the manufacturer's default calibration and is shown for relative comparison among measurement dates; values should only be interpreted with caution as absolute volumetric water content because the sensors were not soil-specifically calibrated.*

| date | start time | end time | nr. of trees measured | air VPD [kPa] | temperature [°C] | relative humidity [%] | wind speed [m/s] | radiation (global) [W/m²] | fraction of diffuse radiation | soil volumetric content [%] at 15 cm | soil volumetric content [%] at 50 cm |
|---|---|---|---|---|---|---|---|---|---|---|---|
| **2024-06-12** | 07:15:28 | 16:05:07 | 15 | 1.21 | 18.5 | 42.8 | 1.0 | 524.5 | 0.5 | 23.0 | 21.3 |
| **2024-06-13** | 07:28:13 | 15:48:11 | 12 | 1.18 | 18.1 | 43.8 | 0.9 | 632.8 | 0.3 | 22.0 | 21.1 |
| **2024-06-18** | 07:12:36 | 15:36:50 | 16 | 1.64 | 26.0 | 50.5 | 1.0 | 736.9 | 0.3 | 18.5 | 20.3 |
| **2024-06-24** | 06:37:37 | 15:28:34 | 19 | 1.65 | 24.0 | 45.2 | 0.8 | 853.2 | 0.2 | 18.4 | 19.5 |
| **2024-06-25** | 06:24:11 | 15:30:19 | 15 | 2.02 | 26.7 | 43.3 | 1.0 | 811.4 | 0.2 | 17.7 | 19.2 |
| **2024-06-26** | 07:11:17 | 15:38:10 | 16 | 2.60 | 31.1 | 43.6 | 0.9 | 786.3 | 0.3 | 16.8 | 18.8 |
| **2024-06-27** | 07:03:48 | 15:32:28 | 9 | 2.00 | 29.2 | 51.1 | 1.0 | 517.8 | 0.6 | 16.0 | 18.5 |
| **2024-07-09** | 06:17:32 | 15:16:31 | 12 | 2.55 | 30.5 | 40.6 | 0.8 | 727.0 | 0.2 | 16.0 | 17.2 |

*Table S 2: Needle widths and the relative proportion of the LI-600N (LI-COR) chamber length occupied by needles during stomatal conductance measurements for each conifer species. Leaf width was entered into the LI-600N configuration as the species-specific rounded mean width, based on measurements of 10 needles per species. The proportion of chamber length occupied by the needle was visually estimated prior to the stomatal conductance measurements and used to recalculate the effective leaf area in the output data. (Please note: The measured mean needle width of* Larix decidua *was below the manufacturer's recommended minimum width of 1 mm for the LI-600N.)*

| Species | Leaf width (mm) | Fraction of chamber length crossed (%) |
|---|---|---|
| **Abi_alb** | 1.8 | 80 |
| **Abi_gra** | 2.4 | 100 |
| **Ced_deo** | 1.1 | 100 |
| **Lar_dec** | 0.6 | 70 |
| **Pic_abi** | 1.2 | 50 |
| **Pic_sit** | 1.4 | 50 |
| **Pin_nig** | 1.4 | 100 |
| **Pin_pon** | 1.6 | 100 |
| **Pin_syl** | 1.4 | 100 |
| **Pse_men** | 1.4 | 60 |
| **Tax_bac** | 2.2 | 50 |

*Table S 3: Prior specifications for the Bayesian stomatal conductance model. For the lognormal distribution, μ and σ represent the mean and standard deviation of the variable's natural logarithm, respectively. For the student-t distribution, df represents the degrees of freedom (governing tail heaviness), μ is the location parameter and σ is the scale parameter. For the Gamma distribution α represents the shape parameter and β represents the rate parameter. Bounds (lower, upper) truncate the distributions to restrict parameters to biologically plausible ranges.*

| Parameter | Argument | Prior Distribution | Bounds |
|---|---|---|---|
| **Parameter a** | Population coefficients | Lognormal (μ = log(0.7), σ = 0.7) | Lower bound = 0 |
| **Parameter b** | Population coefficients | Lognormal (μ = log(1), σ = 0.7) | Lower bound = 0 |
| **Parameter k** | Population coefficients | Lognormal (μ = 0, σ = 1) | Lower bound = 0, Upper bound = 4 |
| **Residual SD (σ)** | Student-t-scale | Student-t (df = 3, μ = 0, σ= 0.7) | Lower bound = 0 |
| **Degrees of freedom (v)** | Student-t-shape | Gamma (α= 2, β=0.1) | Lower bound = 1 |
| **Group-level SD ($\sigma_{tree}$, $\sigma_{leaf}$)** | Random intercepts | Student-t (df = 3, μ=0, σ=1) | Lower bound = 0 |

*Table S 4: Posterior estimates (means and 95 % credible intervals) for model parameters. Fixed effects are broken down by species for parameters a, b, k, and the light interaction (a: Qamb_sc). Group-level variations (random effect standard deviations for tree_id and tree_id:leaf_id), residual standard deviation (σ), and the distributional shape parameter (v) are shown at the bottom. The overall model fit yielded a Bayesian conditional $R^2$ of 0.87 [95% CI: 0.86, 0.87].*

| Effect_Type | Species | a | a:Qamb_sc | b | k |
|---|---|---|---|---|---|
| **Fixed** | Abi_alb | 0.34 [0.16, 0.57] | 0.09 [0.05, 0.16] | 1.31 [0.83, 1.8] | 1.73 [0.68, 2.89] |
| **Fixed** | Abi_gra | 0.53 [0.26, 0.89] | 0.13 [0.06, 0.24] | 1.76 [1.24, 2.32] | 1.86 [0.84, 2.89] |
| **Fixed** | Ace_neg | 0.30 [0.14, 0.49] | 0.05 [0.03, 0.09] | 0.82 [0.46, 1.23] | 0.93 [0.21, 1.87] |
| **Fixed** | Ace_pla | 0.41 [0.20, 0.65] | 0.13 [0.06, 0.22] | 1.00 [0.61, 1.46] | 0.61 [0.11, 1.49] |
| **Fixed** | Ace_pse | 0.44 [0.24, 0.66] | 0.04 [0.02, 0.06] | 0.75 [0.45, 1.08] | 0.84 [0.23, 1.56] |
| **Fixed** | Aes_hip | 0.39 [0.20, 0.60] | 0.09 [0.06, 0.13] | 0.60 [0.34, 0.88] | 0.41 [0.09, 0.92] |
| **Fixed** | Aln_glu | 0.47 [0.25, 0.71] | 0.14 [0.09, 0.21] | 0.84 [0.52, 1.17] | 0.38 [0.08, 0.89] |
| **Fixed** | Bet_pen | 0.61 [0.39, 0.84] | 0.10 [0.07, 0.14] | 0.54 [0.29, 0.81] | 0.28 [0.06, 0.67] |
| **Fixed** | Car_bet | 0.29 [0.13, 0.48] | 0.04 [0.02, 0.08] | 1.04 [0.64, 1.45] | 1.04 [0.27, 1.96] |
| **Fixed** | Cas_sat | 0.92 [0.67, 1.17] | 0.14 [0.09, 0.19] | 0.67 [0.39, 0.97] | 0.92 [0.31, 1.60] |
| **Fixed** | Ced_deo | 0.61 [0.33, 0.94] | 0.22 [0.11, 0.36] | 1.03 [0.67, 1.42] | 0.32 [0.07, 0.80] |
| **Fixed** | Cor_col | 0.37 [0.19, 0.57] | 0.05 [0.03, 0.08] | 0.87 [0.56, 1.18] | 0.78 [0.21, 1.46] |
| **Fixed** | Cor_dom | 0.85 [0.61, 1.09] | 0.17 [0.13, 0.22] | 0.89 [0.61, 1.17] | 1.36 [0.66, 2.09] |
| **Fixed** | Fag_syl | 0.44 [0.24, 0.66] | 0.04 [0.02, 0.07] | 0.79 [0.46, 1.16] | 1.13 [0.38, 1.94] |
| **Fixed** | Fra_exc | 0.92 [0.66, 1.18] | 0.10 [0.07, 0.14] | 0.65 [0.37, 0.93] | 0.63 [0.16, 1.25] |
| **Fixed** | Jug_reg | 0.39 [0.20, 0.59] | 0.04 [0.02, 0.06] | 0.65 [0.38, 0.94] | 1.17 [0.48, 1.87] |
| **Fixed** | Lar_dec | 0.45 [0.24, 0.69] | 0.14 [0.08, 0.23] | 0.94 [0.56, 1.32] | 0.47 [0.09, 1.20] |
| **Fixed** | Pic_abi | 0.55 [0.27, 0.94] | 0.11 [0.05, 0.21] | 1.38 [0.86, 2.04] | 1.00 [0.17, 2.30] |
| **Fixed** | Pic_sit | 0.49 [0.22, 0.94] | 0.10 [0.04, 0.20] | 1.86 [1.15, 2.72] | 1.97 [0.55, 3.60] |
| **Fixed** | Pin_nig | 0.34 [0.17, 0.55] | 0.06 [0.03, 0.10] | 0.81 [0.47, 1.16] | 0.40 [0.09, 0.95] |
| **Fixed** | Pin_pon | 0.62 [0.38, 0.90] | 0.12 [0.07, 0.19] | 1.20 [0.85, 1.56] | 1.21 [0.47, 2.00] |

| | | | | | |
|---|---|---|---|---|---|
| **Fixed** | Pin_syl | 0.54 [0.30, 0.82] | 0.16 [0.10, 0.23] | 0.95 [0.58, 1.34] | 0.61 [0.15, 1.31] |
| **Fixed** | Pla_his | 0.79 [0.53, 1.08] | 0.10 [0.06, 0.15] | 0.69 [0.39, 1.01] | 0.60 [0.14, 1.21] |
| **Fixed** | Pop_tre | 0.73 [0.47, 0.99] | 0.10 [0.06, 0.15] | 0.57 [0.29, 0.87] | 0.35 [0.08, 0.79] |
| **Fixed** | Pru_avi | 0.89 [0.64, 1.17] | 0.16 [0.12, 0.21] | 0.79 [0.50, 1.10] | 0.79 [0.21, 1.50] |
| **Fixed** | Pse_men | 0.34 [0.15, 0.59] | 0.15 [0.06, 0.27] | 1.58 [1.06, 2.16] | 1.55 [0.43, 2.83] |
| **Fixed** | Que_pub | 0.49 [0.28, 0.71] | 0.12 [0.08, 0.17] | 0.34 [0.15, 0.56] | 0.39 [0.08, 0.88] |
| **Fixed** | Que_rob | 1.74 [1.34, 2.16] | 0.26 [0.16, 0.36] | 1.28 [0.97, 1.62] | 1.81 [1.08, 2.55] |
| **Fixed** | Rob_pse | 0.57 [0.33, 0.85] | 0.10 [0.06, 0.16] | 0.99 [0.64, 1.34] | 1.02 [0.33, 1.77] |
| **Fixed** | Sal_cap | 0.87 [0.63, 1.11] | 0.13 [0.10, 0.17] | 0.45 [0.24, 0.69] | 0.26 [0.06, 0.60] |
| **Fixed** | Sal_fra | 1.34 [1.06, 1.65] | 0.17 [0.11, 0.23] | 0.86 [0.58, 1.15] | 0.56 [0.14, 1.14] |
| **Fixed** | Sor_auc | 0.67 [0.43, 0.92] | 0.07 [0.04, 0.10] | 1.10 [0.80, 1.42] | 1.71 [0.96, 2.47] |
| **Fixed** | Tax_bac | 0.41 [0.20, 0.68] | 0.11 [0.05, 0.19] | 1.40 [0.85, 2.04] | 1.73 [0.47, 3.14] |
| **Fixed** | Til_cor | 0.62 [0.39, 0.86] | 0.10 [0.06, 0.14] | 0.85 [0.56, 1.15] | 1.00 [0.36, 1.67] |
| **Fixed** | Til_tom | 0.44 [0.23, 0.66] | 0.05 [0.03, 0.07] | 0.70 [0.36, 1.06] | 0.96 [0.28, 1.76] |
| **Fixed** | Tor_gla | 0.90 [0.66, 1.15] | 0.24 [0.19, 0.30] | 0.69 [0.43, 0.96] | 0.66 [0.18, 1.28] |
| **Fixed** | Ulm_lae | 0.70 [0.43, 1.02] | 0.16 [0.09, 0.24] | 1.08 [0.73, 1.43] | 1.15 [0.45, 1.90] |
| **Fixed** | Ulm_min | 0.41 [0.21, 0.62] | 0.04 [0.03, 0.07] | 0.63 [0.33, 0.96] | 0.72 [0.18, 1.44] |
| **Random: tree_id** | | 0.18 [0.14, 0.22] | | 0.19 [0.14, 0.26] | 0.52 [0.41, 0.66] |
| **Random: tree_id:leaf_id** | | 0.12 [0.10, 0.13] | | 0.06 [0.04, 0.07] | 0.10 [0.04, 0.14] |
| **Residual (sigma)** | | 0.03 [0.03, 0.03] | | | |
| **Distribution Parameter (nu)** | | 1.89 [1.75, 2.06] | | | |

*Table S 5: Markov Chain Monte Carlo (MCMC) convergence and sampling efficiency diagnostics for all structural model parameters. Cells display the potential scale reduction factor ($\hat{R}$), followed by the bulk and tail effective sample sizes for population-level (fixed) effects and hierarchical variance parameters. Across the entire model architecture – including all individuals tree- and leaf-level random deviations – convergence was very good with a global maximum $\hat{R}$ of 1.01 and a global minimum Bulk-ESS of 343, indicating high stability across all posterior distributions. The overall model fit yielded a Bayesian conditional $R^2$ of 0.87 [95% CI: 0.86, 0.87].*

| Species | Effect Type | $\hat{R}$ a | $\hat{R}$ a: Qamb | $\hat{R}$ b | $\hat{R}$ k | Bulk ESS a | Bulk EES a: Qamb | Bulk ESS b | Bulk ESS k | Tail ESS a | Tail ESS a: Qamb | Tail ESS b | Tail ESS k |
|---|---|---|---|---|---|---|---|---|---|---|---|---|---|
| **Abi_alb** | Fixed | 1.00 | 1.00 | 1.00 | 1.00 | 3347 | 4848 | 3237 | 3590 | 4693 | 4663 | 4250 | 4676 |
| **Abi_gra** | Fixed | 1.00 | 1.00 | 1.00 | 1.00 | 2875 | 4859 | 2903 | 3605 | 4297 | 5125 | 4278 | 4425 |
| **Ace_neg** | Fixed | 1.00 | 1.00 | 1.00 | 1.00 | 3327 | 6786 | 4576 | 5123 | 4382 | 5590 | 5131 | 5081 |
| **Ace_pla** | Fixed | 1.00 | 1.00 | 1.00 | 1.00 | 3609 | 6800 | 4473 | 6830 | 4111 | 5469 | 4777 | 6014 |
| **Ace_pse** | Fixed | 1.00 | 1.00 | 1.00 | 1.00 | 3547 | 8585 | 4830 | 3624 | 5059 | 5587 | 4730 | 4878 |
| **Aes_hip** | Fixed | 1.00 | 1.00 | 1.00 | 1.00 | 3400 | 6958 | 4399 | 5274 | 4216 | 6199 | 4656 | 5600 |
| **Aln_glu** | Fixed | 1.00 | 1.00 | 1.00 | 1.00 | 3303 | 4689 | 3702 | 5629 | 4585 | 5824 | 4586 | 5568 |
| **Bet_pen** | Fixed | 1.00 | 1.00 | 1.00 | 1.00 | 3587 | 7075 | 4518 | 6665 | 4386 | 5612 | 4316 | 5578 |
| **Car_bet** | Fixed | 1.00 | 1.00 | 1.00 | 1.00 | 3387 | 6868 | 3273 | 4533 | 4979 | 6132 | 4274 | 5221 |
| **Cas_sat** | Fixed | 1.00 | 1.00 | 1.00 | 1.00 | 3525 | 5275 | 3651 | 3862 | 4556 | 6276 | 3733 | 4390 |
| **Ced_deo** | Fixed | 1.00 | 1.00 | 1.00 | 1.00 | 3488 | 5312 | 3468 | 7815 | 4689 | 5848 | 4699 | 5945 |
| **Cor_col** | Fixed | 1.00 | 1.00 | 1.00 | 1.00 | 3544 | 6901 | 3836 | 3432 | 4428 | 5846 | 3980 | 5414 |
| **Cor_dom** | Fixed | 1.00 | 1.00 | 1.00 | 1.00 | 3599 | 7258 | 3999 | 3382 | 4717 | 6356 | 4157 | 3896 |
| **Fag_syl** | Fixed | 1.00 | 1.00 | 1.00 | 1.00 | 3667 | 7414 | 4311 | 3511 | 4868 | 6173 | 4639 | 4500 |

| | | | | | | | | | | | | | |
|---|---|---|---|---|---|---|---|---|---|---|---|---|---|
| **Fra_exc** | Fixed | 1.00 | 1.00 | 1.00 | 1.00 | 3987 | 6021 | 3954 | 3622 | 4703 | 5865 | 4435 | 4173 |
| **Jug_reg** | Fixed | 1.00 | 1.00 | 1.00 | 1.00 | 3377 | 9548 | 4449 | 3779 | 4067 | 6278 | 4837 | 4587 |
| **Lar_dec** | Fixed | 1.00 | 1.00 | 1.00 | 1.00 | 3452 | 5564 | 4020 | 7290 | 3873 | 5914 | 4301 | 6121 |
| **Pic_abi** | Fixed | 1.00 | 1.00 | 1.00 | 1.00 | 3567 | 6637 | 2780 | 3630 | 4330 | 6202 | 3921 | 5290 |
| **Pic_sit** | Fixed | 1.00 | 1.00 | 1.00 | 1.00 | 2529 | 5309 | 2109 | 2630 | 3653 | 5309 | 3065 | 2991 |
| **Pin_nig** | Fixed | 1.00 | 1.00 | 1.00 | 1.00 | 3353 | 6575 | 4370 | 7552 | 4611 | 5864 | 4953 | 5575 |
| **Pin_pon** | Fixed | 1.00 | 1.00 | 1.00 | 1.00 | 3738 | 7013 | 4606 | 3809 | 4856 | 5952 | 4875 | 4308 |
| **Pin_syl** | Fixed | 1.00 | 1.00 | 1.00 | 1.00 | 3071 | 4991 | 3374 | 4486 | 3996 | 5928 | 3351 | 5332 |
| **Pla_his** | Fixed | 1.00 | 1.00 | 1.00 | 1.00 | 3451 | 5907 | 4050 | 4380 | 4564 | 5457 | 4118 | 5261 |
| **Pop_tre** | Fixed | 1.00 | 1.00 | 1.00 | 1.00 | 3275 | 5435 | 3906 | 5691 | 4053 | 5889 | 5213 | 5312 |
| **Pru_avi** | Fixed | 1.00 | 1.00 | 1.00 | 1.00 | 4295 | 6997 | 2885 | 2376 | 5166 | 6058 | 4011 | 4226 |
| **Pse_men** | Fixed | 1.00 | 1.00 | 1.00 | 1.00 | 3587 | 4571 | 2923 | 4169 | 4224 | 5363 | 4611 | 5197 |
| **Que_pub** | Fixed | 1.00 | 1.00 | 1.00 | 1.00 | 3443 | 5412 | 5482 | 5727 | 4304 | 5288 | 6105 | 5606 |
| **Que_rob** | Fixed | 1.00 | 1.00 | 1.00 | 1.00 | 3097 | 3953 | 3032 | 2827 | 4014 | 4854 | 4117 | 3835 |
| **Rob_pse** | Fixed | 1.00 | 1.00 | 1.00 | 1.00 | 3242 | 6838 | 3838 | 3749 | 3953 | 6085 | 4333 | 4280 |
| **Sal_cap** | Fixed | 1.00 | 1.00 | 1.00 | 1.00 | 3840 | 7349 | 3893 | 4623 | 4632 | 5936 | 4610 | 4417 |
| **Sal_fra** | Fixed | 1.00 | 1.00 | 1.00 | 1.00 | 3385 | 8010 | 3270 | 3571 | 5263 | 4830 | 4040 | 4297 |
| **Sor_auc** | Fixed | 1.00 | 1.00 | 1.00 | 1.00 | 4011 | 8342 | 4028 | 3784 | 4926 | 5488 | 4511 | 4664 |
| **Tax_bac** | Fixed | 1.00 | 1.00 | 1.00 | 1.00 | 3711 | 5787 | 3147 | 3610 | 4333 | 5234 | 4441 | 4104 |
| **Til_cor** | Fixed | 1.00 | 1.00 | 1.00 | 1.00 | 3598 | 6076 | 4525 | 3853 | 4938 | 5963 | 5258 | 4451 |
| **Til_tom** | Fixed | 1.00 | 1.00 | 1.00 | 1.00 | 3461 | 6535 | 4462 | 3805 | 4283 | 6185 | 4536 | 4859 |
| **Tor_gla** | Fixed | 1.00 | 1.00 | 1.00 | 1.00 | 3678 | 6294 | 3813 | 3730 | 4353 | 6110 | 4829 | 4653 |
| **Ulm_lae** | Fixed | 1.00 | 1.00 | 1.00 | 1.00 | 3324 | 4664 | 3590 | 3717 | 4267 | 6357 | 4629 | 4440 |
| **Ulm_min** | Fixed | 1.00 | 1.00 | 1.00 | 1.00 | 3116 | 6380 | 3830 | 3942 | 4139 | 5315 | 5038 | 4963 |
| **Random: tree_id** | | 1.00 | | 1.00 | 1.00 | 2594 | | 2020 | 2228 | 2594 | | 2020 | 2228 |
| **Random: tree_id:leaf_id** | | 1.00 | | 1.01 | 1.01 | 1277 | | 343 | 356 | 1277 | | 343 | 356 |
| **Residual (sigma)** | | Rhat =1.00, Sigma Bulk ESS = 3088 | | | | | | | | | | | |
| **Distribution Parameter (nu)** | | Rhat =1.00, Nu Bulk ESS = 4740 | | | | | | | | | | | |

*Table S 6: Model-derived stomatal control metrics across 38 studied species. Values represent the posterior means calculated across all 4,000 posterior draws from the population-level response curves, reconstructed at a discrete 1000-point vector resolution of leaf-to-air vapor pressure deficit ($VPD_L$). Numbers inside the brackets indicate the 95 % Bayesian credible intervals (2.5$^{th}$ and 97.5$^{th}$ percentiles). The maximum operational stomatal conductance to water vapor ($g_{max}$) is identified as the peak value of the curve and does not reflect absolute physiological limits, but operational maximums captured under ambient environmental field conditions. The optimum vapor pressure deficit ($VPD_{opt}$) corresponds to the coordinate where $g_{max}$ is achieved. The critical vapor pressure deficit threshold ($VPD_{50}$) defines the point along the post-peak declining limb where stomatal conductance drops to exactly 50 % of $g_{max}$. The maximum relative rate of stomatal closure ($S_{closure}$) is determined by evaluating the steepest negative first derivative along the post-peak smoothed declining limb of the relative conductance curve (standardized by $g_{max}$).*

| **Species** | **$g_{max}$ (mol m$^{-2}$ s$^{-1}$)** | **$VPD_{opt}$ (kPa)** | **$VPD_{50}$ (kPa)** | **$S_{closure}$ (mol m$^{-2}$ s$^{-1}$ kPa$^{-1}$)** |
|---|---|---|---|---|
| **Abi_alb** | 0.106 [0.048, 0.192] | 1.32 [0.66, 1.97] | 2.90 [1.97, 3.99] | 0.41 [0.51, 0.29] |
| **Abi_gra** | 0.094 [0.052, 0.153] | 1.06 [0.60, 1.52] | 2.26 [1.61, 2.96] | 0.53 [0.66, 0.41] |
| **Ace_neg** | 0.145 [0.062, 0.258] | 1.13 [0.33, 2.06] | 3.18 [1.80, 4.97] | 0.31 [0.44, 0.19] |
| **Ace_pla** | 0.181 [0.085, 0.327] | 0.59 [0.14, 1.23] | 2.02 [1.14, 3.18] | 0.44 [0.63, 0.28] |
| **Ace_pse** | 0.226 [0.120, 0.378] | 1.13 [0.35, 2.15] | 3.29 [1.80, 5.42] | 0.30 [0.44, 0.18] |
| **Aes_hip** | 0.237 [0.119, 0.386] | 0.71 [0.15, 1.70] | 2.90 [1.52, 5.21] | 0.29 [0.45, 0.16] |

| | | | | |
|---|---|---|---|---|
| **Aln_glu** | 0.255 [0.133, 0.422] | 0.46 [0.11, 1.05] | 1.96 [1.10, 3.26] | 0.42 [0.61, 0.26] |
| **Bet_pen** | 0.403 [0.245, 0.586] | 0.55 [0.12, 1.40] | 2.78 [1.49, 5.21] | 0.29 [0.45, 0.15] |
| **Car_bet** | 0.112 [0.047, 0.202] | 1.00 [0.30, 1.74] | 2.65 [1.53, 3.99] | 0.39 [0.53, 0.26] |
| **Cas_sat** | 0.532 [0.344, 0.884] | 1.41 [0.48, 2.61] | 3.92 [2.16, 6.56] | 0.26 [0.38, 0.15] |
| **Ced_deo** | 0.324 [0.173, 0.550] | 0.31 [0.08, 0.77] | 1.48 [0.87, 2.45] | 0.53 [0.77, 0.34] |
| **Cor_col** | 0.167 [0.079, 0.280] | 0.90 [0.27, 1.68] | 2.70 [1.53, 4.20] | 0.35 [0.50, 0.23] |
| **Fag_syl** | 0.229 [0.116, 0.412] | 1.44 [0.59, 2.43] | 3.71 [2.25, 5.72] | 0.28 [0.40, 0.18] |
| **Fra_exc** | 0.515 [0.355, 0.739] | 1.00 [0.26, 2.09] | 3.31 [1.73, 5.69] | 0.28 [0.42, 0.16] |
| **Jug_reg** | 0.265 [0.117, 0.555] | 1.85 [0.78, 3.23] | 4.66 [2.75, 7.55] | 0.23 [0.33, 0.14] |
| **Lar_dec** | 0.219 [0.106, 0.384] | 0.50 [0.11, 1.14] | 1.93 [1.07, 3.15] | 0.44 [0.65, 0.27] |
| **Pic_abi** | 0.165 [0.082, 0.320] | 0.69 [0.17, 1.31] | 1.90 [1.05, 2.79] | 0.52 [0.70, 0.37] |
| **Pic_sit** | 0.082 [0.040, 0.154] | 1.03 [0.42, 1.53] | 2.19 [1.41, 2.91] | 0.55 [0.70, 0.41] |
| **Pin_nig** | 0.183 [0.083, 0.322] | 0.50 [0.12, 1.14] | 2.10 [1.16, 3.56] | 0.40 [0.60, 0.23] |
| **Pin_pon** | 0.200 [0.121, 0.307] | 1.01 [0.45, 1.62] | 2.52 [1.64, 3.56] | 0.42 [0.55, 0.31] |
| **Pin_syl** | 0.239 [0.133, 0.384] | 0.64 [0.17, 1.29] | 2.17 [1.22, 3.45] | 0.41 [0.59, 0.26] |
| **Pla_his** | 0.426 [0.284, 0.621] | 0.89 [0.23, 1.89] | 3.01 [1.61, 5.24] | 0.30 [0.45, 0.18] |
| **Pop_tre** | 0.456 [0.299, 0.643] | 0.64 [0.15, 1.58] | 2.91 [1.53, 5.51] | 0.29 [0.45, 0.15] |
| **Pru_avi** | 0.434 [0.290, 0.640] | 1.01 [0.32, 1.88] | 3.00 [1.70, 4.65] | 0.32 [0.46, 0.21] |
| **Pse_men** | 0.077 [0.034, 0.147] | 0.97 [0.33, 1.55] | 2.21 [1.26, 3.09] | 0.52 [0.69, 0.38] |
| **Que_pub** | 0.380 [0.213, 0.608] | 1.27 [0.26, 3.30] | 5.32 [2.45, 11.14] | 0.17 [0.29, 0.07] |
| **Que_rob** | 0.562 [0.380, 0.895] | 1.42 [0.87, 2.03] | 3.06 [2.22, 4.10] | 0.39 [0.49, 0.30] |
| **Rob_pse** | 0.227 [0.135, 0.356] | 1.05 [0.38, 1.77] | 2.78 [1.65, 4.16] | 0.37 [0.50, 0.25] |
| **Sal_cap** | 0.611 [0.427, 0.821] | 0.62 [0.14, 1.55] | 3.27 [1.70, 6.10] | 0.25 [0.39, 0.13] |
| **Sal_fra** | 0.648 [0.461, 0.918] | 0.65 [0.17, 1.34] | 2.28 [1.29, 3.60] | 0.39 [0.55, 0.26] |
| **Sor_auc** | 0.272 [0.156, 0.475] | 1.56 [0.90, 2.27] | 3.42 [2.39, 4.69] | 0.34 [0.44, 0.25] |
| **Cor_dom** | 0.422 [0.257, 0.740] | 1.57 [0.78, 2.48] | 3.71 [2.40, 5.38] | 0.30 [0.40, 0.21] |
| **Tor_gla** | 0.484 [0.331, 0.700] | 0.97 [0.27, 1.94] | 3.12 [1.71, 5.24] | 0.30 [0.43, 0.18] |
| **Tax_bac** | 0.112 [0.054, 0.198] | 1.21 [0.48, 1.82] | 2.68 [1.76, 3.63] | 0.44 [0.56, 0.31] |
| **Til_cor** | 0.284 [0.171, 0.443] | 1.19 [0.44, 2.04] | 3.18 [1.91, 4.91] | 0.32 [0.44, 0.21] |
| **Til_tom** | 0.248 [0.124, 0.448] | 1.41 [0.48, 2.58] | 3.90 [2.19, 6.53] | 0.26 [0.38, 0.15] |
| **Ulm_lae** | 0.252 [0.160, 0.376] | 1.08 [0.47, 1.73] | 2.73 [1.73, 3.96] | 0.39 [0.51, 0.27] |
| **Ulm_min** | 0.235 [0.116, 0.396] | 1.15 [0.30, 2.31] | 3.62 [1.95, 6.37] | 0.26 [0.40, 0.14] |

*Table S 7: Species-level leaf trait means. Mean values are shown with standard deviations in brackets (±SD) for stomatal density (SD), guard cell length (GCL), specific leaf area (SLA), leaf dry matter content (LDMC), leaf thickness (Thick), leaf toughness (Tough), and leaf nitrogen per area (Leaf N (area)).*

| species | SD [no. mm$^{-2}$] | GCL [μm] | SLA [m$^2$ kg$^{-1}$] | LDMC [g g$^{-1}$] | Thick [mm] | Tough [N mm$^{-1}$] | Leaf N (area) [kg m$^{-2}$] |
|---|---|---|---|---|---|---|---|
| **Abi_alb** | 49.35 (±8.31) | 46.26 (±5.93) | 39.29 (±4.48) | 0.444 (±0.034) | 0.659 (±0.067) | 4.84 (±0.40) | 0.0316 (±0.0063) |
| **Abi_gra** | 48.40 (±4.91) | 38.95 (±5.87) | 39.04 (±3.19) | 0.416 (±0.007) | 0.746 (±0.113) | 4.02 (±0.51) | 0.0319 (±0.0011) |
| **Ace_neg** | 616.65 (±214.20) | 18.94 (±2.32) | 157.13 (±16.74) | 0.366 (±0.029) | 0.175 (±0.027) | 1.05 (±0.11) | 0.0082 (±0.0002) |
| **Ace_pla** | 260.82 (±80.24) | 21.35 (±2.74) | 126.16 (±9.82) | 0.447 (±0.027) | 0.161 (±0.009) | 1.18 (±0.07) | 0.0129 (±0.0026) |
| **Ace_pse** | 95.01 (±64.86) | 26.65 (±4.25) | 106.10 (±1.61) | 0.418 (±0.021) | 0.210 (±0.012) | 1.60 (±0.16) | 0.0143 (±0.0042) |
| **Aes_hip** | 335.57 (±85.59) | 22.87 (±3.29) | 113.85 (±14.76) | 0.398 (±0.025) | 0.164 (±0.011) | 0.76 (±0.06) | 0.0086 (±0.0025) |

| | | | | | | | |
|---|---|---|---|---|---|---|---|
| **Aln_glu** | 215.87 (±46.43) | 30.60 (±7.10) | 126.33 (±4.39) | 0.405 (±0.012) | 0.191 (±0.017) | 0.98 (±0.04) | 0.0233 (±0.0007) |
| **Bet_pen** | 155.79 (±80.60) | 32.41 (±5.78) | 124.98 (±13.33) | 0.394 (±0.019) | 0.187 (±0.019) | 1.08 (±0.17) | 0.0212 (±0.0009) |
| **Car_bet** | 233.34 (±34.98) | 26.67 (±3.02) | 126.66 (±24.31) | 0.474 (±0.024) | 0.168 (±0.037) | 0.79 (±0.02) | 0.0155 (±0.0017) |
| **Cas_sat** | 329.05 (±60.64) | 24.75 (±6.31) | 90.38 (±3.93) | 0.413 (±0.029) | 0.209 (±0.010) | 1.24 (±0.23) | 0.0221 (±0.0055) |
| **Ced_deo** | 38.33 (±15.66) | 38.16 (±5.39) | 31.42 (±5.62) | 0.459 (±0.002) | 0.824 (±0.151) | 5.62 (±0.07) | 0.0310 (±0.0067) |
| **Cor_col** | 127.15 (±31.16) | 32.50 (±8.26) | 126.23 (±9.69) | 0.438 (±0.001) | 0.183 (±0.007) | 1.09 (±0.12) | 0.0146 (±0.0023) |
| **Cor_dom** | 161.38 (±34.48) | 31.02 (±3.89) | 97.90 (±5.50) | 0.471 (±0.017) | 0.197 (±0.017) | 1.08 (±0.10) | 0.0144 (±0.0019) |
| **Fag_syl** | 272.00 (±43.62) | 23.49 (±3.64) | 116.84 (±10.37) | 0.487 (±0.005) | 0.177 (±0.019) | 1.29 (±0.04) | 0.0177 (±0.0003) |
| **Fra_exc** | 321.83 (±59.19) | 26.38 (±3.87) | 111.79 (±1.60) | 0.378 (±0.038) | 0.190 (±0.009) | 1.05 (±0.20) | 0.0194 (±0.0031) |
| **Jug_reg** | 295.52 (±43.60) | 30.32 (±5.07) | 128.79 (±13.04) | 0.359 (±0.012) | 0.215 (±0.025) | 1.07 (±0.08) | 0.0150 (±0.0041) |
| **Lar_dec** | 55.15 (±7.98) | 31.03 (±2.97) | 103.33 (±27.06) | 0.383 (±0.126) | 0.216 (±0.045) | 1.54 (±0.31) | 0.0247 (±0.0102) |
| **Pic_abi** | 32.08 (±6.15) | 41.60 (±5.29) | 26.57 (±2.02) | 0.502 (±0.036) | 0.890 (±0.052) | 5.91 (±0.39) | 0.0419 (±0.0028) |
| **Pic_sit** | 47.51 (±10.57) | 34.49 (±5.42) | 38.35 (±6.63) | 0.443 (±0.063) | 0.669 (±0.128) | 5.63 (±1.26) | 0.0289 (±0.0059) |
| **Pin_nig** | 56.46 (±11.37) | 39.58 (±4.37) | 29.37 (±1.62) | 0.454 (±0.040) | 0.865 (±0.069) | 11.27 (±1.00) | 0.0361 (±0.0035) |
| **Pin_pon** | 66.19 (±9.03) | 27.93 (±3.32) | 38.79 (±9.61) | 0.401 (±0.077) | 0.888 (±0.070) | 8.36 (±0.99) | 0.0450 (±0.0094) |
| **Pin_syl** | 81.67 (±12.38) | 26.76 (±5.98) | 32.86 (±4.58) | 0.425 (±0.030) | 0.829 (±0.077) | 12.36 (±1.10) | 0.0409 (±0.0068) |
| **Pla_his** | 183.04 (±46.71) | 42.28 (±5.78) | 111.89 (±11.25) | 0.403 (±0.019) | 0.215 (±0.017) | 1.46 (±0.12) | 0.0135 (±0.0022) |
| **Pop_tre** | 183.50 (±60.11) | 19.42 (±3.42) | 131.36 (±19.56) | 0.450 (±0.005) | 0.208 (±0.042) | 2.03 (±0.54) | 0.0170 (±0.0025) |
| **Pru_avi** | 405.67 (±91.90) | 24.17 (±3.27) | 116.75 (±8.99) | 0.423 (±0.010) | 0.177 (±0.031) | 0.82 (±0.07) | 0.0145 (±0.0025) |
| **Pse_men** | 73.54 (±13.95) | 25.46 (±2.86) | 44.22 (±18.75) | 0.465 (±0.036) | 0.449 (±0.038) | 2.00 (±0.54) | 0.0342 (±0.0158) |
| **Que_pub** | 597.32 (±104.91) | 26.02 (±2.57) | 82.09 (±22.54) | 0.491 (±0.021) | 0.226 (±0.028) | 1.80 (±0.39) | 0.0260 (±0.0047) |
| **Que_rob** | 403.34 (±66.56) | 29.17 (±2.84) | 100.71 (±9.42) | 0.450 (±0.027) | 0.211 (±0.043) | 1.34 (±0.25) | 0.0255 (±0.0008) |
| **Rob_pse** | 339.76 (±116.58) | 16.34 (±2.48) | 198.42 (±25.46) | 0.364 (±0.060) | 0.138 (±0.009) | 0.63 (±0.08) | 0.0164 (±0.0035) |
| **Sal_cap** | 759.87 (±234.70) | 14.03 (±2.44) | 94.37 (±5.04) | 0.419 (±0.006) | 0.234 (±0.019) | 1.19 (±0.06) | 0.0247 (±0.0006) |
| **Sal_fra** | 202.13 (±44.07) | 28.26 (±3.83) | 90.41 (±13.83) | 0.375 (±0.035) | 0.281 (±0.040) | 1.41 (±0.16) | 0.0193 (±0.0038) |
| **Sor_auc** | 117.60 (±23.19) | 30.26 (±4.39) | 119.54 (±10.78) | 0.450 (±0.012) | 0.186 (±0.018) | 0.86 (±0.09) | 0.0097 (±0.0006) |
| **Tax_bac** | 64.50 (±16.21) | 25.52 (±8.62) | 44.24 (±7.37) | 0.445 (±0.048) | 0.548 (±0.067) | 2.70 (±0.45) | 0.0268 (±0.0037) |
| **Til_cor** | 176.05 (±30.56) | 29.61 (±3.36) | 124.14 (±25.21) | 0.404 (±0.009) | 0.199 (±0.025) | 0.97 (±0.07) | 0.0215 (±0.0030) |
| **Til_tom** | 290.16 (±53.43) | 23.68 (±3.49) | 119.40 (±13.27) | 0.436 (±0.024) | 0.191 (±0.020) | 0.81 (±0.12) | 0.0174 (±0.0021) |
| **Tor_gla** | 210.05 (±43.78) | 28.00 (±3.71) | 105.75 (±12.48) | 0.475 (±0.009) | 0.203 (±0.020) | 1.70 (±0.26) | 0.0168 (±0.0033) |
| **Ulm_lae** | 405.20 (±67.57) | 26.35 (±2.69) | 134.66 (±15.43) | 0.423 (±0.005) | 0.145 (±0.014) | 0.80 (±0.14) | 0.0105 (±0.0020) |

| | | | | | | | |
|---|---|---|---|---|---|---|---|
| **Ulm_min** | 367.24 (±64.98) | 36.81 (±6.22) | 106.92 (±15.83) | 0.413 (±0.020) | 0.231 (±0.032) | 0.97 (±0.10) | 0.0154 (±0.0028) |

*Table S 8: Summary of Bayesian linear regression models analysing the interactive effects of plant functional traits and tree group (broadleaved vs. conifer) on stomatal behaviour metrics (N=28 models). Traits (SD, GCL, SLA, LDMC, thickness, toughness, and leaf N per area) were z-score scaled prior to analysis to facilitate comparison. For each model, all posterior draws of the stomatal behaviour metrics were used, thereby propagating uncertainty in the estimated metrics through the regression analysis. For each model, the Bayesian coefficient of determination (Bayes $R^2$), absolute group-specific slopes, and 95% credible intervals (CI) are reported; models where the 95% credible intervals do not cross zero are highlighted in bold. Model convergence and diagnostic performance are assessed via the potential scale reduction factor ($\hat{R}$), as well as Bulk and Tail Effective Sample Sizes (ESS).*

| **Model name** | **response** | **Predictor trait (scaled)** | **Tree group** | **Bayes $R^2$** | **slope** | **CI lower** | **CI upper** | $\hat{R}$ | **ESS Bulk** | **ESS Tail** |
|---|---|---|---|---|---|---|---|---|---|---|
| Model 1 | gmax | SD | broadleaved | 0.27 | 0.022 | -0.04 | 0.08 | 1.00 | 2709 | 2259 |
| | | | conifer | 0.27 | -0.006 | -0.90 | 0.89 | 1.00 | 1734 | 1833 |
| Model 2 | gmax | GCL | broadleaved | 0.25 | 0.006 | -0.05 | 0.07 | 1.00 | 2296 | 1724 |
| | | | conifer | 0.25 | -0.002 | -0.08 | 0.07 | 1.00 | 2348 | 2386 |
| **Model 3** | **gmax** | **SLA** | **broadleaved** | **0.43** | **-0.128** | **-0.21** | **-0.04** | 1.00 | 2198 | 2138 |
| | | | conifer | 0.43 | 0.016 | -0.12 | 0.15 | 1.00 | 2137 | 2103 |
| Model 4 | gmax | LDMC | broadleaved | 0.25 | -0.009 | -0.06 | 0.04 | 1.00 | 3607 | 3012 |
| | | | conifer | 0.25 | -0.015 | -0.10 | 0.08 | 1.00 | 3529 | 2940 |
| **Model 5** | **gmax** | **thick** | **broadleaved** | **0.51** | **0.753** | **0.39** | **1.16** | 1.00 | 1153 | 1305 |
| | | | conifer | 0.51 | 0.027 | -0.04 | 0.10 | 1.00 | 1191 | 1240 |
| **Model 6** | **gmax** | **tough** | **broadleaved** | **0.38** | **0.535** | **0.14** | **0.94** | 1.00 | 833 | 920 |
| | | | conifer | 0.38 | 0.029 | -0.02 | 0.08 | 1.00 | 855 | 1021 |
| **Model 7** | **gmax** | **leaf_N_area** | **broadleaved** | **0.45** | **0.150** | **0.06** | **0.24** | 1.00 | 2069 | 2221 |
| | | | conifer | 0.45 | 0.035 | -0.06 | 0.12 | 1.00 | 2315 | 2165 |
| Model 8 | VPDopt | SD | broadleaved | 0.13 | 0.012 | -0.18 | 0.21 | 1.00 | 2757 | 2115 |
| | | | conifer | 0.13 | 1.065 | -1.50 | 3.53 | 1.00 | 1829 | 1655 |
| Model 9 | VPDopt | GCL | broadleaved | 0.14 | 0.054 | -0.16 | 0.26 | 1.00 | 2293 | 2454 |
| | | | conifer | 0.14 | -0.034 | -0.26 | 0.20 | 1.00 | 2257 | 2275 |
| Model 10 | VPDopt | SLA | broadleaved | 0.13 | -0.026 | -0.35 | 0.31 | 1.00 | 2289 | 2030 |
| | | | conifer | 0.13 | -0.059 | -0.50 | 0.35 | 1.00 | 1648 | 1616 |
| Model 11 | VPDopt | LDMC | broadleaved | 0.17 | 0.103 | -0.08 | 0.28 | 1.00 | 2756 | 2586 |
| | | | conifer | 0.17 | -0.049 | -0.29 | 0.18 | 1.00 | 2940 | 2895 |
| Model 12 | VPDopt | thick | broadleaved | 0.12 | -0.207 | -1.50 | 1.15 | 1.00 | 1269 | 1548 |
| | | | conifer | 0.12 | -0.047 | -0.30 | 0.21 | 1.00 | 1323 | 1713 |
| Model 13 | VPDopt | tough | broadleaved | 0.15 | -0.445 | -2.02 | 1.04 | 1.01 | 1047 | 1066 |
| | | | conifer | 0.15 | -0.071 | -0.26 | 0.11 | 1.00 | 1060 | 1127 |
| Model 14 | VPDopt | leaf_N_area | broadleaved | 0.14 | -0.119 | -0.45 | 0.20 | 1.00 | 2264 | 2261 |
| | | | conifer | 0.14 | 0.007 | -0.31 | 0.34 | 1.00 | 2310 | 2134 |
| Model 15 | VPD50 | SD | broadleaved | 0.27 | 0.114 | -0.28 | 0.51 | 1.00 | 2626 | 2297 |
| | | | conifer | 0.27 | 2.210 | -1.40 | 6.06 | 1.00 | 1646 | 1940 |
| Model 16 | VPD50 | GCL | broadleaved | 0.25 | 0.067 | -0.36 | 0.48 | 1.00 | 2181 | 2164 |
| | | | conifer | 0.25 | -0.054 | -0.39 | 0.29 | 1.00 | 2113 | 2157 |
| Model 17 | VPD50 | SLA | broadleaved | 0.26 | -0.168 | -0.71 | 0.38 | 1.00 | 2186 | 2263 |
| | | | conifer | 0.26 | 0.004 | -0.68 | 0.72 | 1.00 | 1915 | 1762 |
| Model 18 | VPD50 | LDMC | broadleaved | 0.26 | 0.094 | -0.23 | 0.42 | 1.00 | 2377 | 2631 |
| | | | conifer | 0.26 | -0.116 | -0.47 | 0.25 | 1.00 | 2340 | 2550 |
| Model 19 | VPD50 | thick | broadleaved | 0.25 | 0.416 | -2.04 | 2.81 | 1.00 | 924 | 995 |
| | | | conifer | 0.25 | -0.063 | -0.48 | 0.36 | 1.00 | 969 | 1022 |
| Model 20 | VPD50 | tough | broadleaved | 0.25 | 0.242 | -2.54 | 3.07 | 1.00 | 945 | 1474 |
| | | | conifer | 0.25 | -0.021 | -0.33 | 0.28 | 1.00 | 961 | 1446 |
| Model 21 | VPD50 | leaf_N_area | broadleaved | 0.24 | -0.094 | -0.68 | 0.50 | 1.00 | 2145 | 2125 |
| | | | conifer | 0.24 | 0.012 | -0.48 | 0.51 | 1.00 | 2223 | 1742 |
| Model 22 | Sclosure | SD | broadleaved | 0.62 | -0.022 | -0.05 | 0.01 | 1.00 | 2322 | 2203 |
| | | | conifer | 0.62 | -0.364 | -0.98 | 0.26 | 1.00 | 1803 | 2090 |
| Model 23 | Sclosure | GCL | broadleaved | 0.56 | 0.002 | -0.03 | 0.04 | 1.00 | 2465 | 2098 |
| | | | conifer | 0.56 | 0.004 | -0.04 | 0.05 | 1.00 | 2287 | 2464 |
| Model 24 | Sclosure | SLA | broadleaved | 0.59 | 0.037 | -0.01 | 0.08 | 1.00 | 2856 | 2667 |
| | | | conifer | 0.59 | -0.014 | -0.13 | 0.11 | 1.00 | 1743 | 2315 |
| Model 25 | Sclosure | LDMC | broadleaved | 0.58 | -0.001 | -0.03 | 0.02 | 1.00 | 2933 | 2322 |
| | | | conifer | 0.58 | 0.022 | -0.04 | 0.08 | 1.00 | 2586 | 2629 |
| Model 26 | Sclosure | thick | broadleaved | 0.60 | -0.203 | -0.43 | 0.03 | 1.00 | 1172 | 1417 |
| | | | conifer | 0.60 | -0.003 | -0.07 | 0.06 | 1.00 | 1256 | 1692 |

| | | | | | | | | | | |
|---|---|---|---|---|---|---|---|---|---|---|
| Model 27 | Sclosure | tough | broadleaved | 0.61 | -0.174 | -0.38 | 0.05 | 1.01 | 949 | 1302 |
| | | | conifer | 0.61 | -0.019 | -0.06 | 0.02 | 1.01 | 1008 | 1301 |
| Model 28 | Sclosure | leaf_N_area | broadleaved | 0.58 | -0.025 | -0.07 | 0.02 | 1.00 | 2718 | 2440 |
| | | | conifer | 0.58 | -0.016 | -0.08 | 0.05 | 1.00 | 2493 | 2357 |

*Table S 9: Summary of Bayesian linear regression models analysing the interactive effects of leaf carbon isotope discrimination (Δδ13C) and tree group (broadleaved/conifer) on stomatal behaviour metrics. Leaf Δδ13C was z-score scaled prior to analysis. For each model the Bayesian coefficient of determination (Bayes $R^2$), absolute group-specific slopes, and 95 % credible intervals (CI) are reported; models where the 95 % credible interval do not cross zero are highlighted in bold. Model convergence and diagnostic performance are assessed via the potential scale reduction factor ($\hat{R}$), as well as Bulk and Tail Effective Sample Sizes (ESS).*

| **Model** | **Response** | **Predictor** | $\hat{R}$ | **ESS Bulk** | **ESS Tail** | **Slope Broad-leaved** | **CI Broad-leaved** | **Slope conifer** | **CI conifer** | **Bayes $R^2$** |
|---|---|---|---|---|---|---|---|---|---|---|
| **gmax_ Dd13C** | Dd13C | gmax | 1.00 | 3089 | 2911 | 0.009 | [-0.041, 0.058] | 0.054 | [-0.018, 0.126] | 0.272 |
| **VPDopt_ Dd13C** | Dd13C | VPDopt | 1.00 | 4446 | 4824 | **-0.174** | **[-0.308, -0.034]** | **-0.220** | **[-0.36, -0.081]** | 0.341 |
| **VPD50_ Dd13C** | Dd13C | VPD50 | 1.00 | 2411 | 2426 | -0.226 | [-0.496, 0.057] | **-0.268** | **[-0.52, -0.006]** | 0.340 |
| **Sclosure_ Dd13C** | Dd13C | Sclosure | 0.99 | 2580 | 2410 | 0.009 | [-0.019, 0.038] | 0.011 | [-0.041, 0.062] | 0.575 |

#### Supplementary Figures

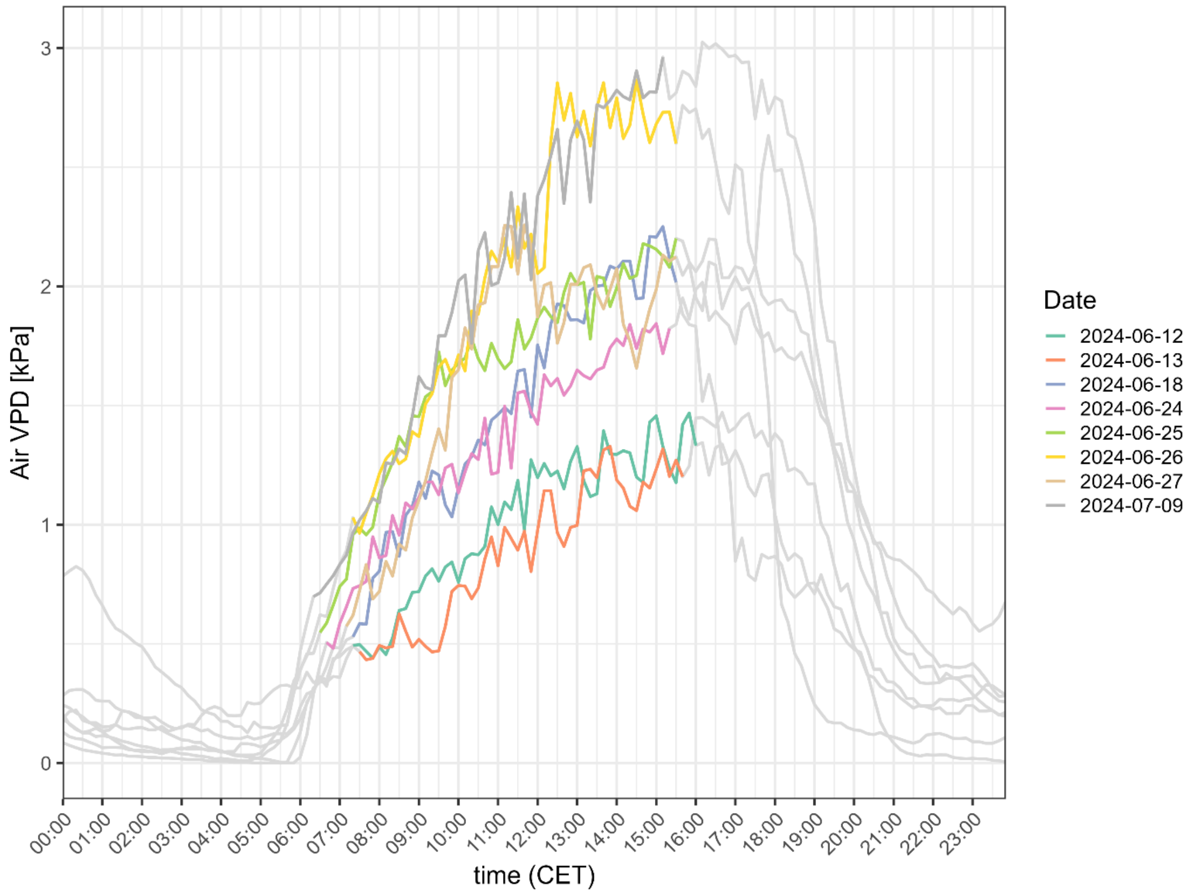

*Figure S 1: Diurnal VPD dynamics. Air vapour pressure deficit (VPD; kPa) across eight measurement days, plotted against time (CET). Measurements were obtained from an on-site climate station (WXT520, Vaisala, Finland), recording at 10-min intervals. Lines are coloured by date during the measurement period of stomatal conductance, while values outside this period are shown in grey. VPD was derived from air temperature and relative humidity as VPD = sVP − aVP, where saturation vapour pressure (sVP) was calculated following Magnus-Tetens' approximation (Tetens, 1930).*

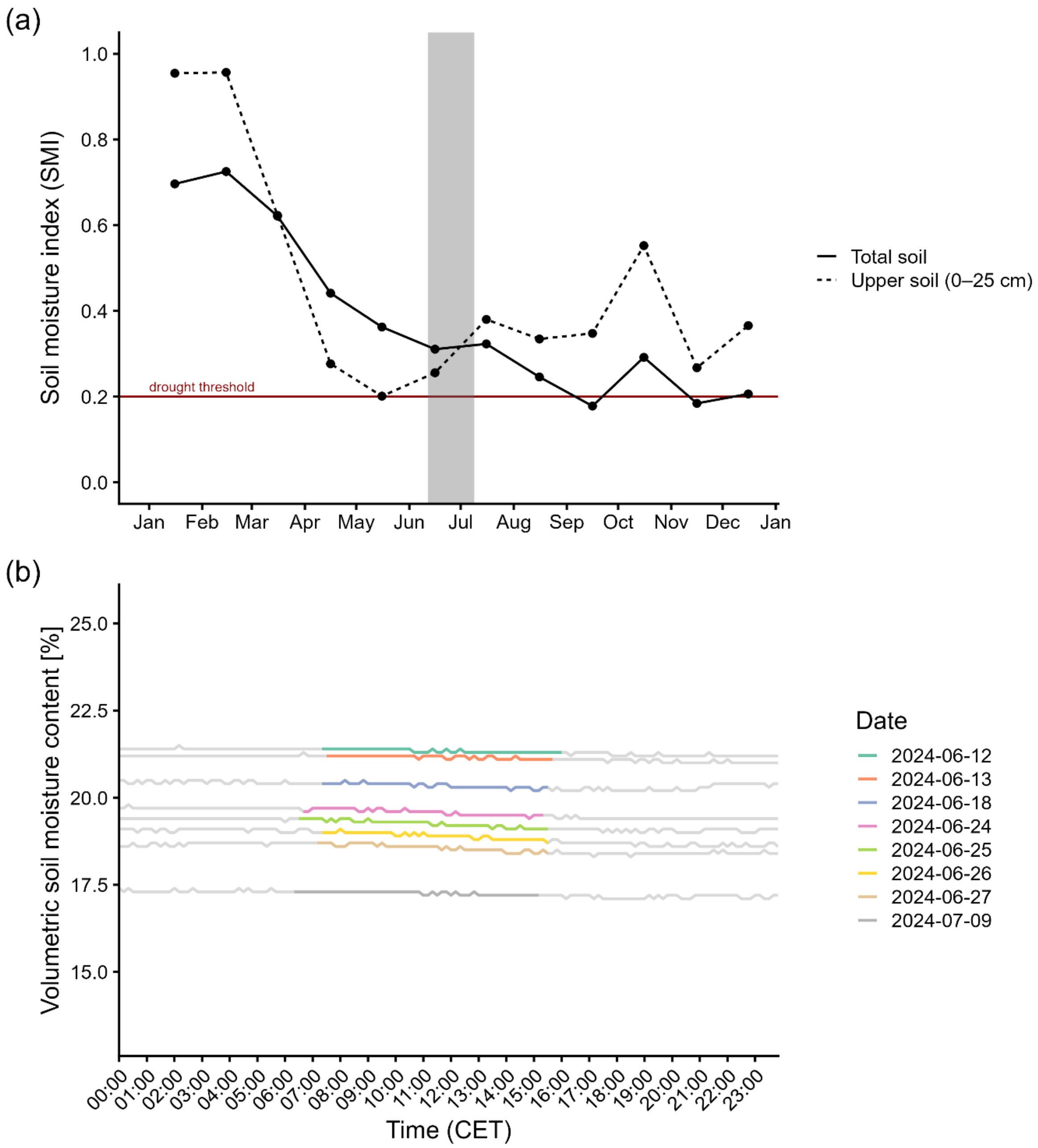

*Figure S 2: Soil moisture status. (a) Monthly soil moisture index (SMI) in 2024 at the ARBOfun site, extracted from the gridded Historical Drought Monitor v2 dataset (Boeing et al., 2022), provided by the UFZ at a spatial resolution of 0.015625° (approximately 1.2 km; Drought Monitor Germany (https://www.ufz.de/index.php?en=37937). SMI values are scaled from 0 to 1 and standardized relative to the 1974–2023 reference period, with lower values indicating drier conditions. The horizontal line indicates the drought threshold (SMI = 0.20), corresponding to the 20th percentile of the reference distribution. SMI is shown for the upper soil layer (0–25 cm) and the total soil profile. (b) Volumetric soil moisture content [%] at 50 cm depth across eight measurement days, plotted against Central European Time (CET). Lines are coloured by date during the measurement period of stomatal conductance, while values outside this period are shown in grey. Measurements were obtained from an on-site ML3 ThetaProbe soil moisture sensor (Delta-T Devices, UK), recording at 10-minute intervals. Soil water content was measured using the manufacturer's default calibration and is shown for relative comparison among measurement dates; values should only be interpreted with caution as absolute volumetric water content because the sensors were not soil-specifically calibrated.*

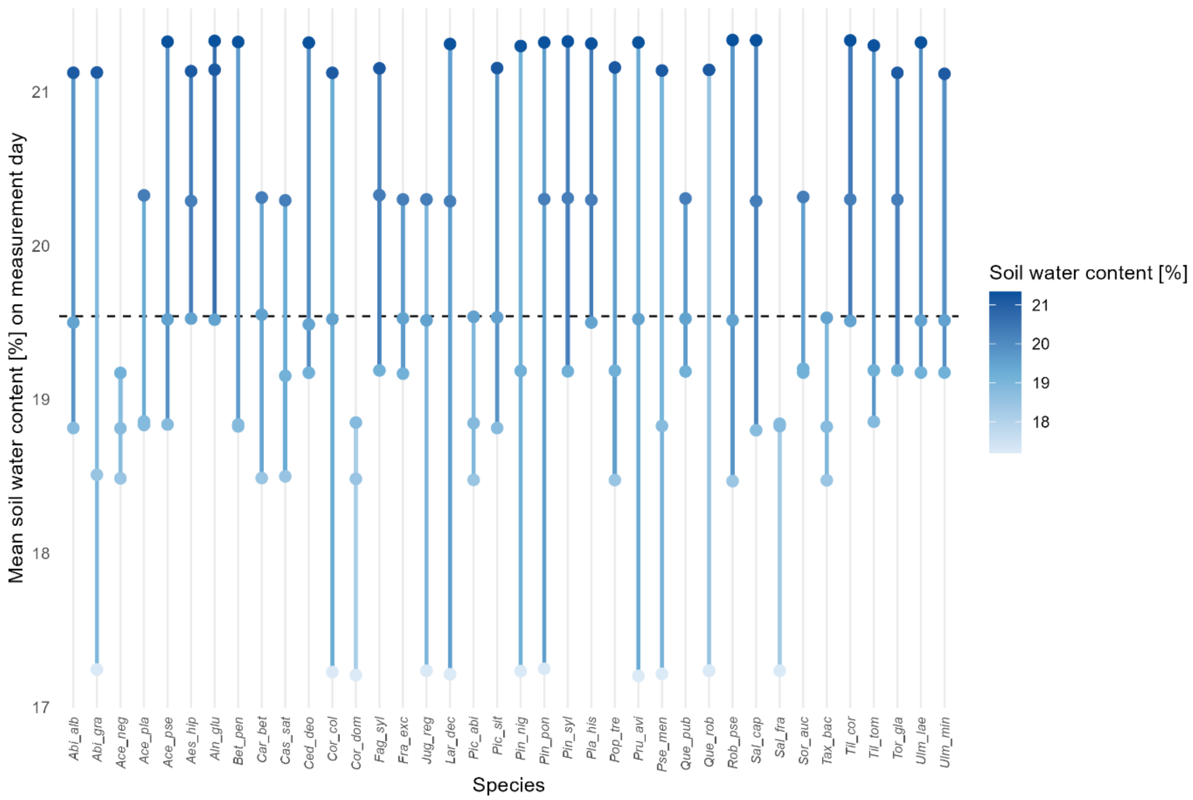

*Figure S 3: Soil water content across measurement days and species. Mean soil water content [%] measured on each of the three sampling days is shown for 38 tree species, with three individual measurements per species (points) representing the trees measured on each day. In most cases, the three trees of a given species were measured on three different days. Points are connected by lines, indicating species-specific range of soil water content across the three measurement days. Horizontal reference line indicates the overall mean soil water content across all species and measurement days. Measurements were obtained from an on-site ML3 ThetaProbe soil moisture sensor (Delta-T Devices, UK), recording at 10-minute intervals. Soil water content was measured using the manufacturer's default calibration and is shown for relative comparison among measurement dates; values should only be interpreted with caution as absolute volumetric water content because the sensors were not soil-specifically calibrated.*

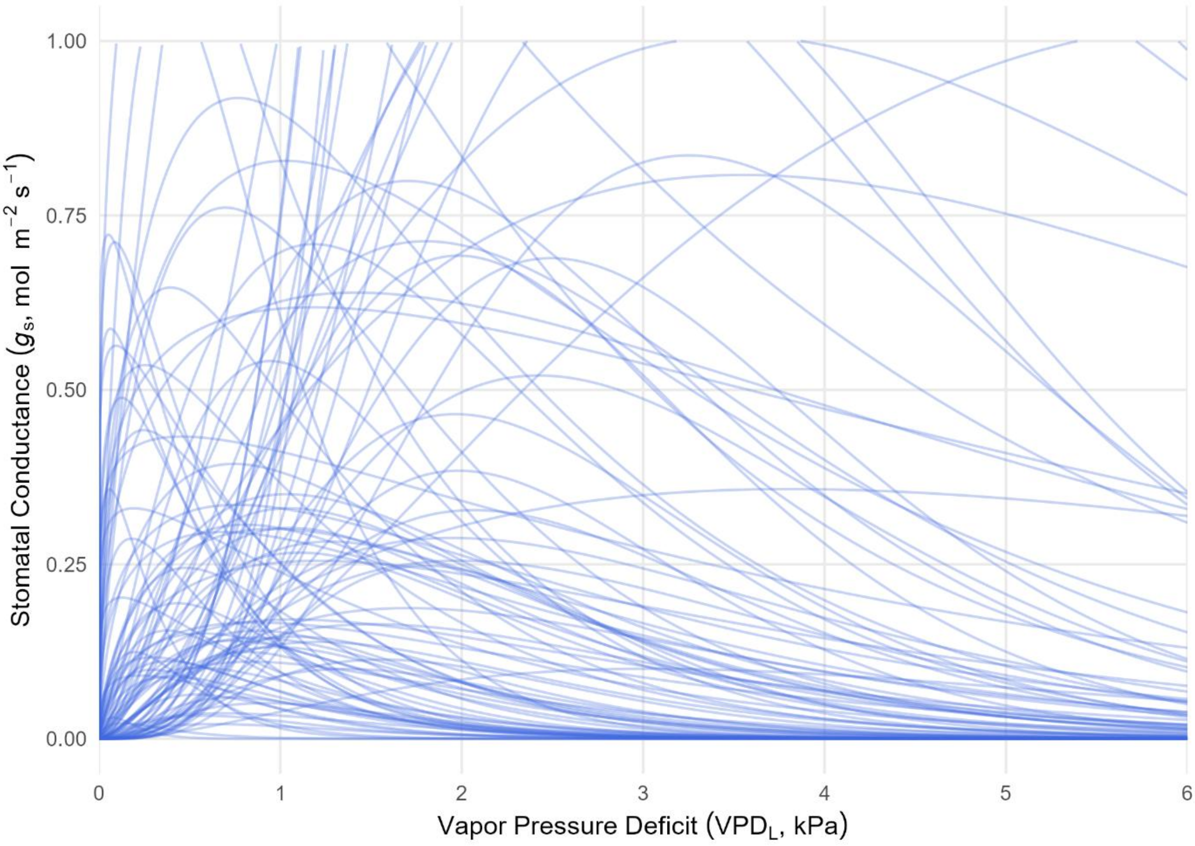

*Figure S 4: Prior predictive check for the non-linear Bayesian stomatal conductance model. The plot displays 100 random curves simulated entirely from the joint prior distribution across a range of $VPD_L$ (holding $Q_{amb}$ at their mean level). The simulated curves consistently fall within biologically realistic boundaries, demonstrating that the chosen priors are regularizing and scientifically plausible without overly restricting potential data spaces.*

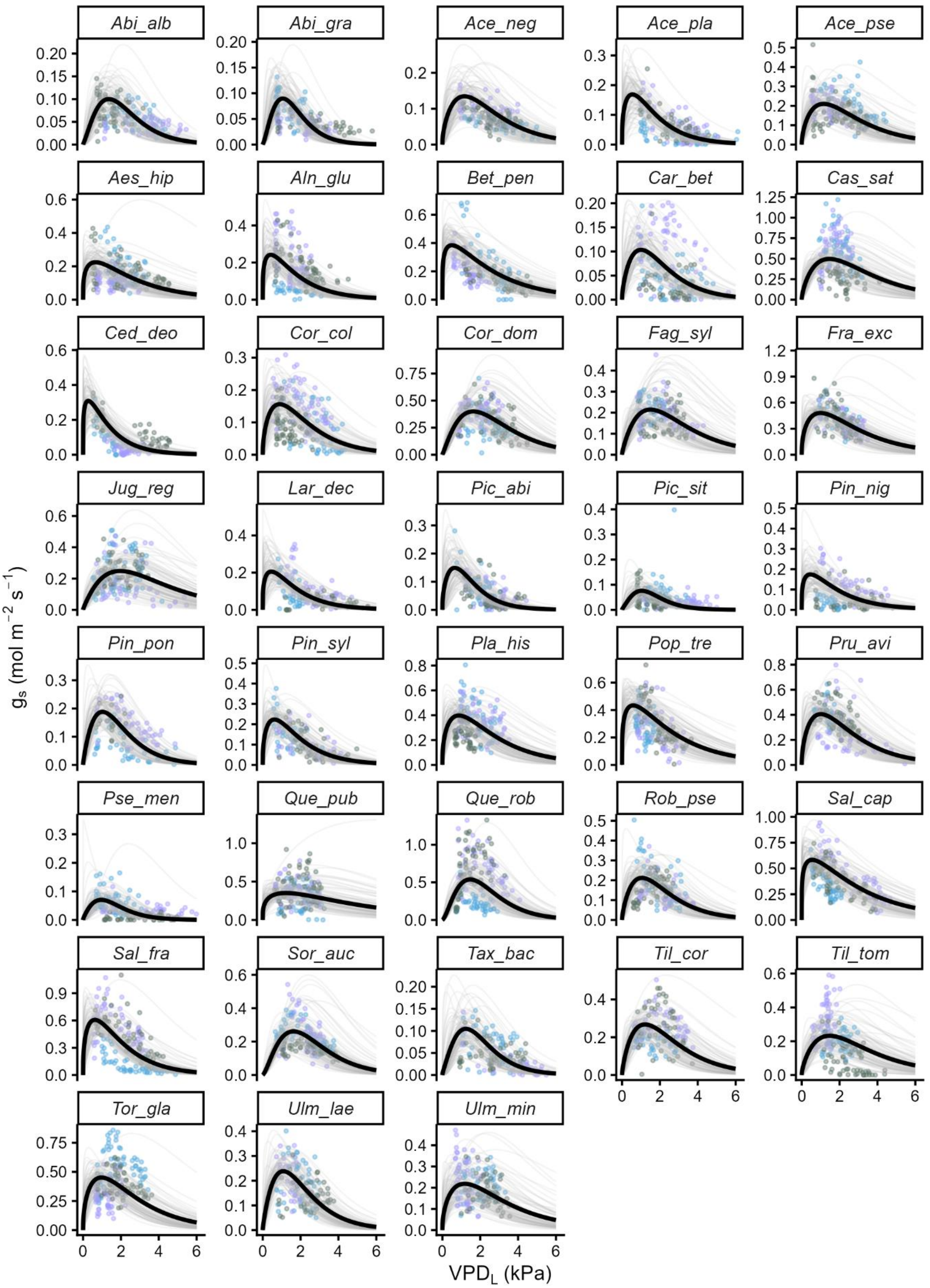

*Figure S 5: Species-specific responses of stomatal conductance ($g_{sw}$) to leaf-to-air vapor pressure deficit ($VPD_L$). Black curves represent expected population-level mean responses calculated via posterior predictions from the non-linear Bayesian mixed-effects model, holding ambient light ($Q_{amb}$) constant at the mean level. Thin grey lines represent a selection of 100 random posterior draws per species. Datapoints are the raw measurement data, coloured by tree individual (3 trees per species). See Table 1 for species abbreviations.*

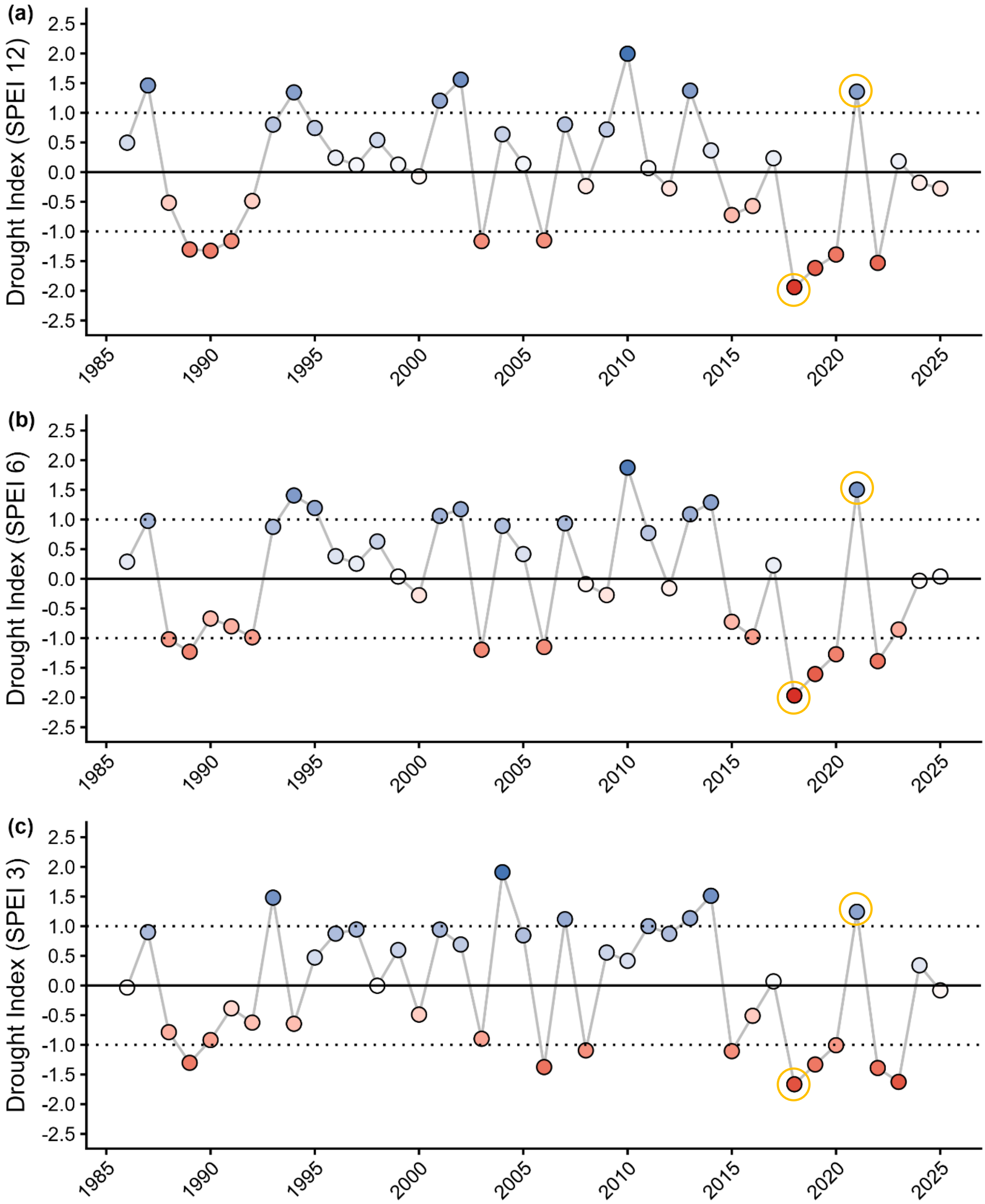

*Figure S 6: Annual Standardized Precipitation–Evapotranspiration Index (SPEI) for the study region calculated for (a) the full year (12-month SPEI), (b) the growing season (April–September; 6-month SPEI), and (c) the summer season (June–August; 3-month SPEI) from 1985 to 2025. Negative SPEI values indicate drier-than-average conditions, whereas positive values indicate wetter-than-average conditions relative to the long-term climatic mean. Horizontal dashed lines indicate SPEI values of ±1, corresponding to commonly used thresholds for drought (≤ −1) and wet (≥ 1) conditions. The years relevant for this study are highlighted with the yellow circle, the dry year 2018 and the wet year 2021. For further information on the calculation of SPEI, see Vincente-Serrano et al. (2010) and Beguería et al. (2014).*

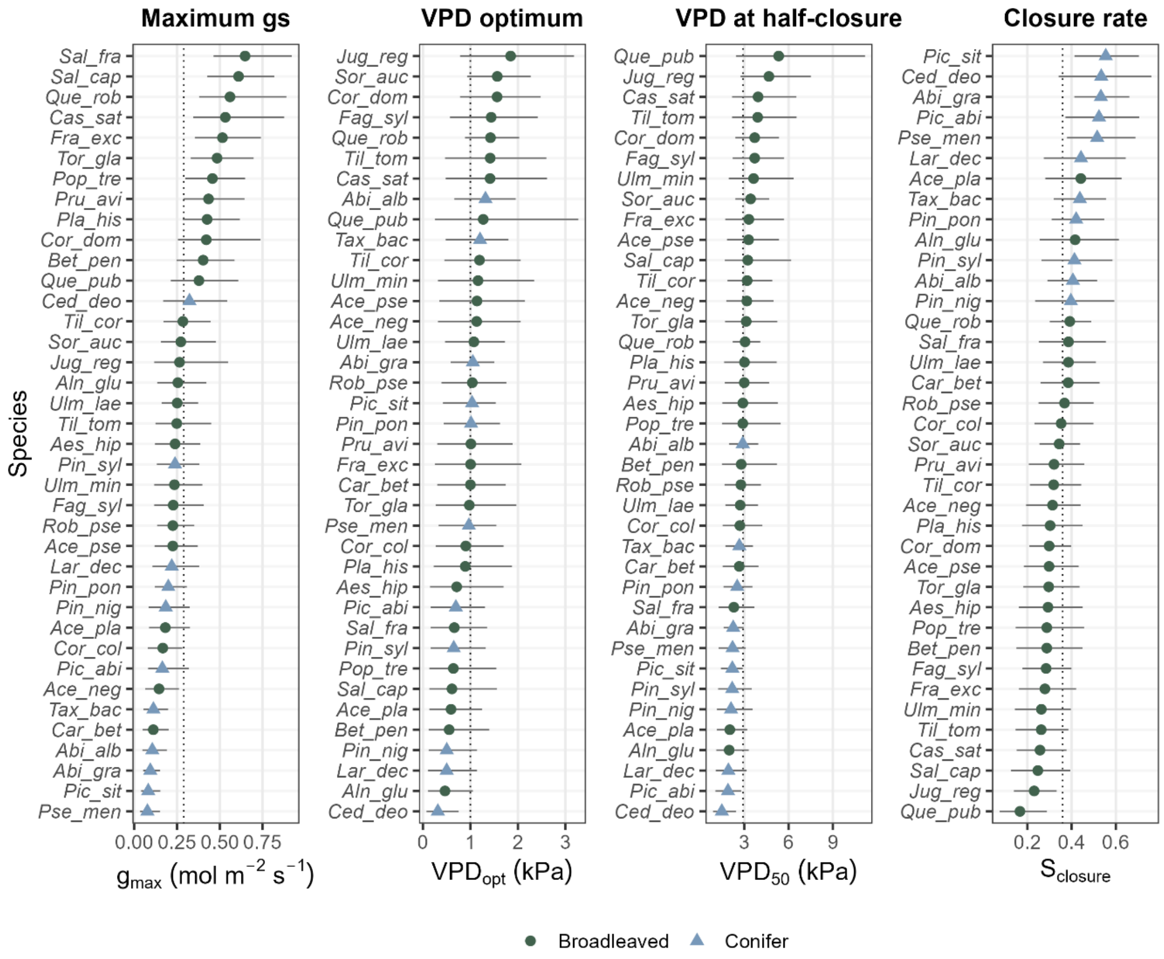

*Figure S 7: Stomatal behaviour metrics. Species-level estimates of maximum stomatal conductance ($g_{max}$), VPD optimum ($VPD_{opt}$), VPD at half-closure ($VPD_{50}$) and the maximum rate of stomatal closure ($S_{closure}$). Points show posterior mean estimates and horizontal lines represent 95 % credible intervals. The dotted line represents the across-species mean for each metric. Broadleaved and conifer species are distinguished by symbol and colour.*

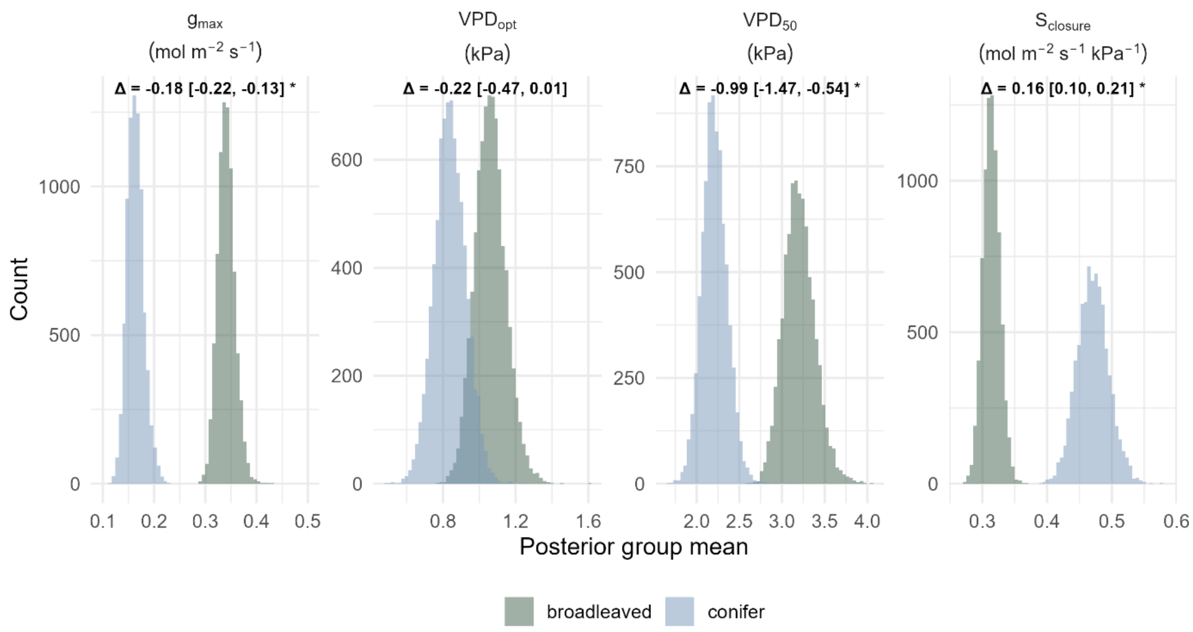

*Figure S 8: Differences in stomatal behaviour metrics between broadleaved and conifer species. Distributions show posterior estimates of clade-level means for each stomatal behaviour metric. Δ denotes the posterior mean difference between conifers and broadleaved species, with 95% credible intervals shown in brackets. Asterisks indicate 95% credible intervals that do not overlap zero.*

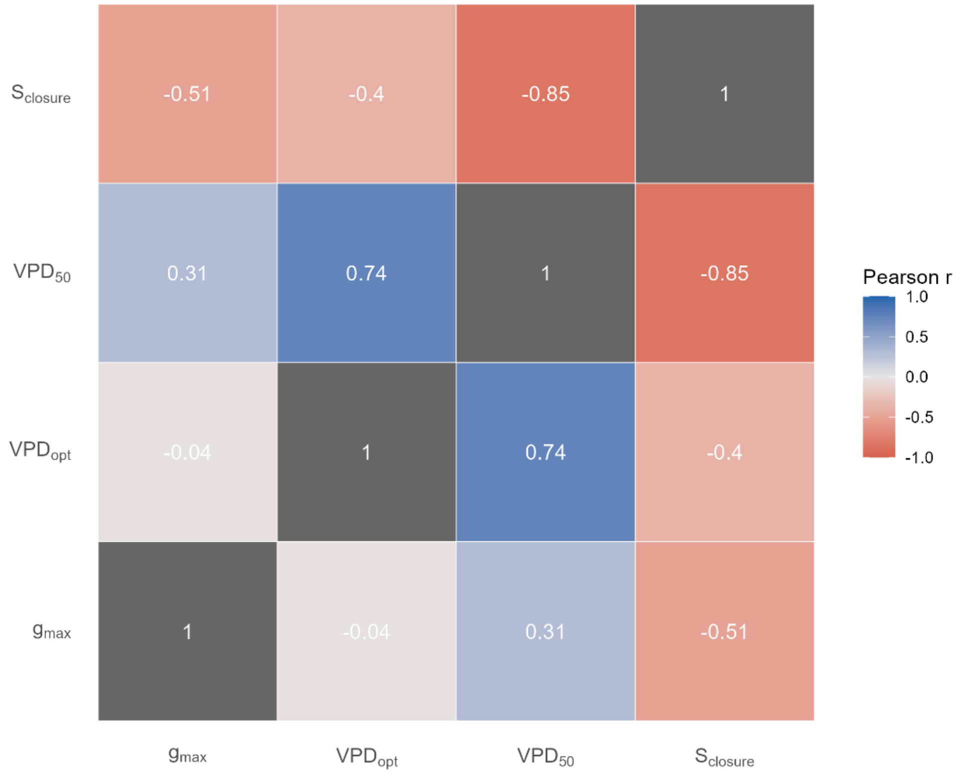

*Figure S 9: Pearson correlation matrix of the four stomatal behaviour metrics: maximum stomatal conductance ($g_{max}$), VPD optimum ($VPD_{opt}$), VPD at half closure ($VPD_{50}$) and the stomatal closure rate ($S_{closure}$). Colour intensity reflects the strength and direction of the correlation (blue = positive, red = negative).*

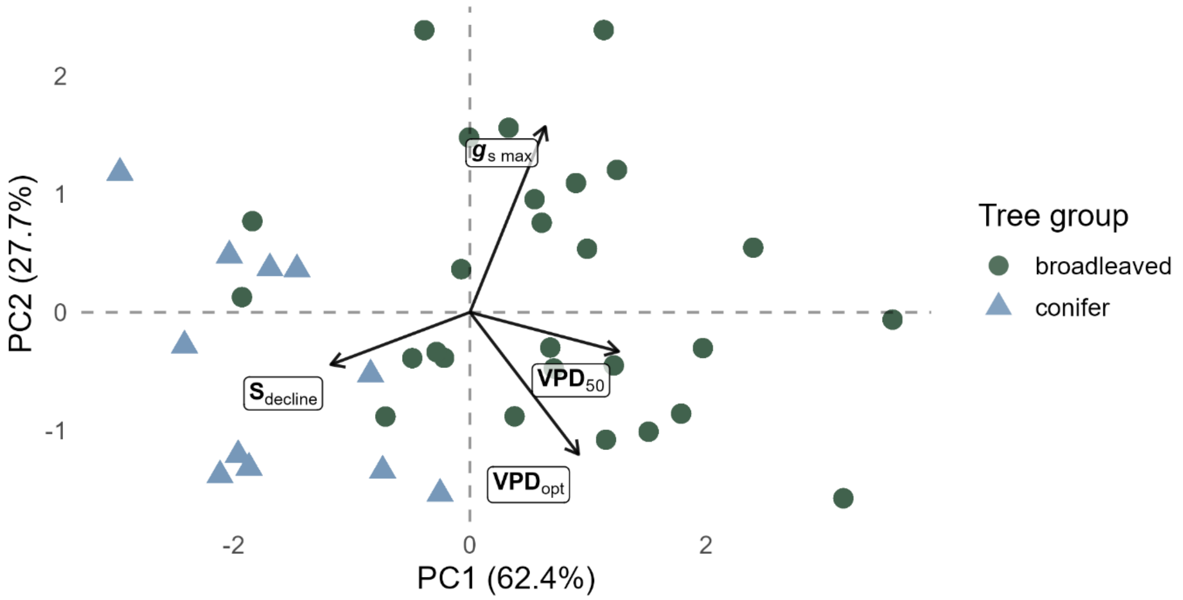

*Figure S 10: Principal Component Analysis (PCA) of stomatal behaviour metrics. The PCA is based on four metrics describing stomatal behaviour: maximum stomatal conductance ($g_{max}$), the optimum VPD ($VPD_{opt}$), the VPD at which $g_s$ is reduced by 50 % of its maximum ($VPD_{50}$) and the maximal slope of the conductance decline ($S_{closure}$). Points represent species positioned in the multivariate space. Arrows indicate the direction and strength of loadings on the first two principal component axes.*

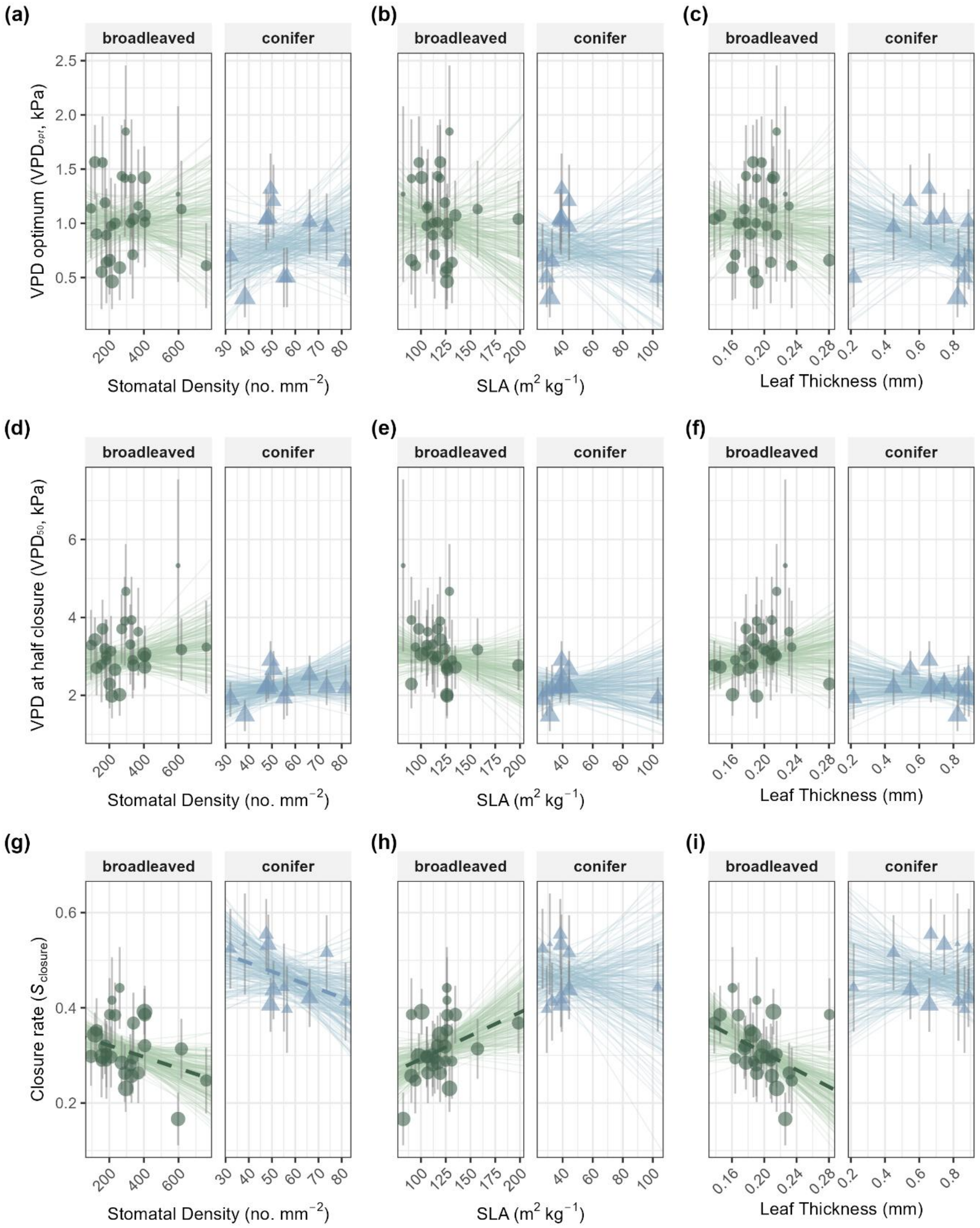

*Figure S 11: Relationship between stomatal sensitivity metrics ($VPD_{opt}$, $VPD_{50}$, and $S_{closure}$), and selected leaf functional traits. Models included an interaction between tree group and leaf trait; groups are shown in separate panels because of their different trait ranges. Individual panels show (a,d,g) Stomatal density (SD), (b,e,h) Specific Leaf Area (SLA), (c,f,i) Leaf Thickness.* Larix decidua*, the only deciduous conifer in our species selection, was an outlier in some models due to its exceptionally high SLA (103.3 $m^2$ $kg^{-1}$) and very thin needles (0.22 mm). No relationship was supported by a 95% credible interval excluding zero. Dashed lines indicate marginal support (66% credible interval). Thin lines illustrate the posterior uncertainty cloud based on 200 random draws from the model parameters. Error bars on the points indicate the standard deviation (±SD) around individual species means, and point size is scaled inversely with variance (1/SD) to reflect parameter precision in model weighting. Bayes $R^2$ of the models are: (a) 0.13, (b) 0.13, (c) 0.12, (d) 0.27, (e) 0.26, (f) 0.25, (g) 0.62, (h) 0.59, (i) 0.60. Please note that $R^2$ values reflect total model fit, which in some cases is driven primarily by intercept differences between tree groups (conifer/broadleaved) rather than by credible slope relationships with the main predictor. See Table S9 for detailed model summaries.*

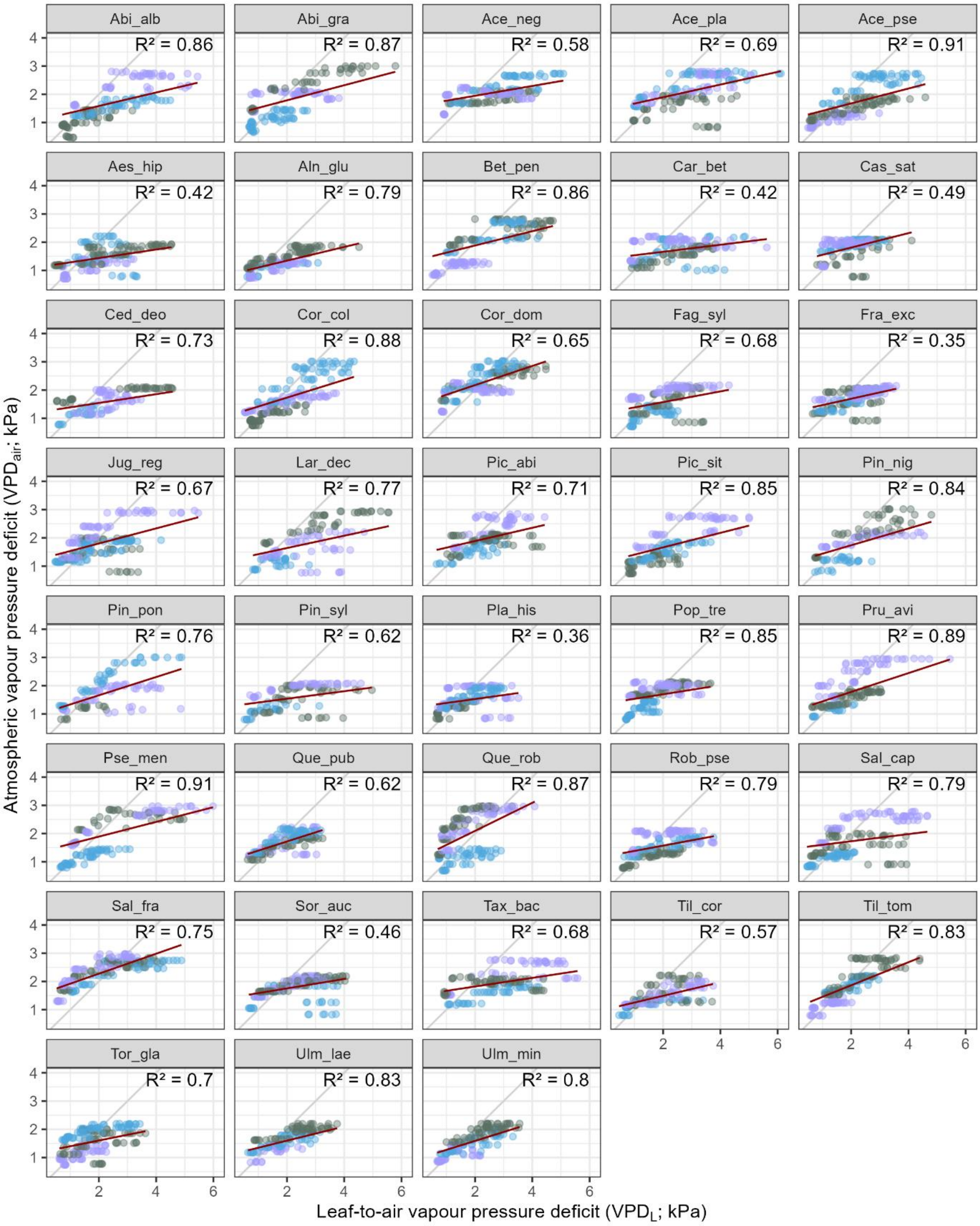

*Figure S 12: Species-specific relationship between ambient VPD (y axis) and leaf-to-air VPD (x-axis). For ambient VPD, measurements were obtained from an on-site climate station (WXT520, Vasiala, Finland), recording at 10-min intervals. Ambient VPD was derived from air temperature and relative humidity as VPD = sVP − aVP, where saturation vapour pressure (sVP) was calculated following Magnus-Tetens' approximation(Tetens, 1930). It was interpolated to the nearest measurement in time of leaf-to-air VPD ($VPD_L$) measured by the porometer devices (LI-600, LI-600N, LICOR). Point colours indicate the three different tree individuals per species. Grey line indicates the 1:1 relationship. Regression lines were obtained from simple linear models. For interpretation: The greater the deviation of the regression line from the 1:1 line, the greater the difference between leaf-to-air and ambient VPD, reflecting stronger leaf warming or cooling relative to air temperature.*

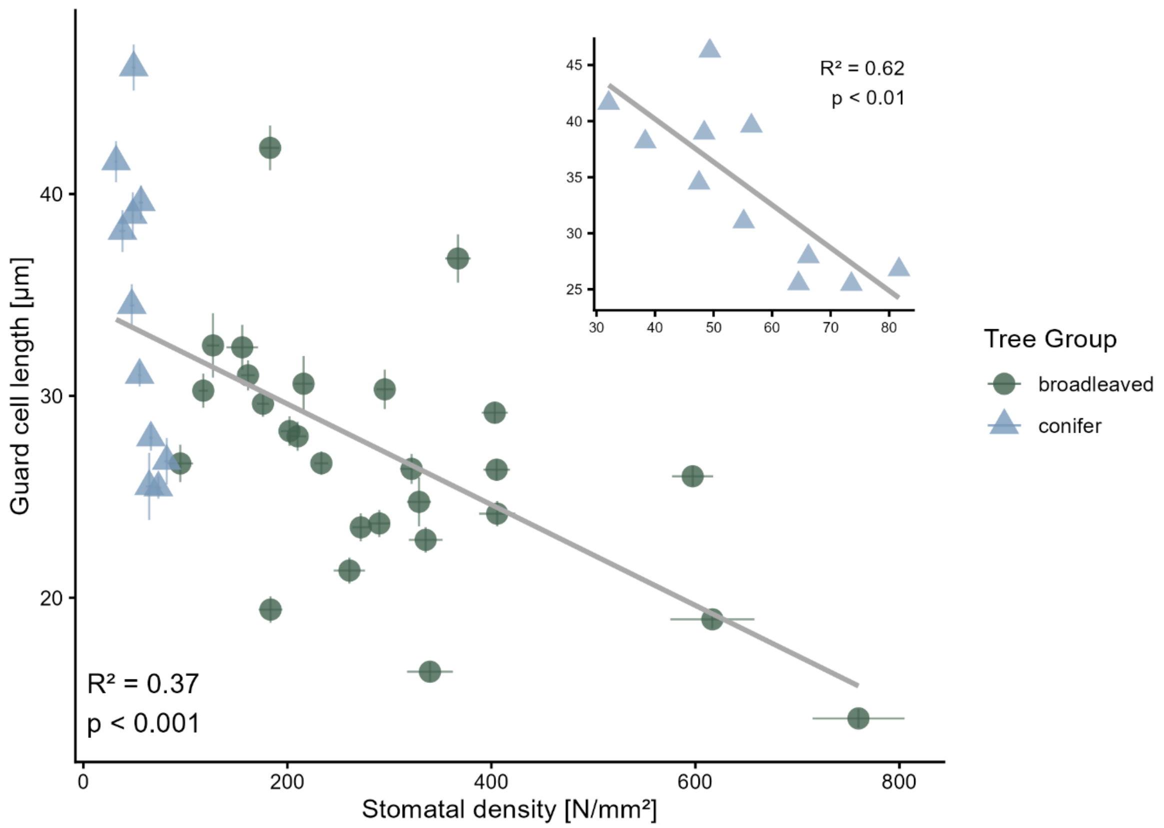

*Figure S 13: Relationship between stomatal density and guard cell length across species. Species means are shown with horizontal and vertical error bars indicating ± standard error for stomatal density and guard cell length, respectively. Point colour and shape distinguish broadleaved species from conifers. The solid line represents the linear regression fitted across all species. The inset shows the corresponding relationship for conifers only.*
